# Understanding the physical processes behind DNA-DNA proximity ligation assays

**DOI:** 10.1101/2025.05.02.649190

**Authors:** Bernardo J. Zubillaga Herrera, Amit Das, Linden Burack, Ailung Wang, Michele Di Pierro

## Abstract

In the last decade, DNA-DNA proximity ligation assays opened powerful new ways to study the 3D organization of genomes and have become a mainstay experimental technology. Yet many aspects of these experiments remain poorly understood. We study the inner workings of DNA-DNA proximity ligation assays through numerical experiments and theoretical modeling. Chromosomes are modeled at nucleosome resolution and evolved in time via molecular dynamics. A virtual Hi-C experiment reproduces, in-silico, the different steps of the Hi-C protocol, including: crosslinking of chromatin to an underlying proteic matrix, enzymatic digestion of DNA, and subsequent proximity ligation of DNA open ends. The protocol is simulated on ensembles of different structures as well as individual structures, enabling the construction of ligation maps and the calculation of ligation probabilities as functions of genomic and Euclidean distance. The methods help to assess the effect of the many variables of the Hi-C experiment and of subsequent data processing methods on the quality of the final results.

## 1. INTRODUCTION

Experimental studies on 3D chromatin organization reveal the complexity of large-scale arrangements of DNA in nuclei, thought to correlate with macromolecular machineries of transcription and regulation, DNA replication and repair, and overall genome functionality (e.g. loop extrusion or enhancer-promoter contacts). Chromosome conformation capture (3C) techniques are molecular biology experiments that probe chromatin’s spatial organization by quantifying interaction frequencies (contacts) between loci in 3D physical proximity, although possibly separated by large genomic distances. ^1–10^

Hi-C, a successful 3C technology, quantifies interactions between all possible pairs of DNA fragments at a given resolution. Hi-C’s power for chromosome architecture capture stems from comprehensive detection through proximity ligation and deep sequencing. Ligation maps reveal a wealth of complexity through checkerboard patterns, and structures like domains, loops, and compartments, harnessing the high throughput of Hi-C (processing billions of pair-end sequence reads) ^1–7,11–26^.

Ensemble Hi-C measures 3D contact frequency between fragments pairs, averaging over cell ensembles, aggregating proximity-contact reads into ligation maps. Single-cell Hi-C (scHi-C) probes individual cells, suggesting that chromosomes in single cells exhibit domain structure, at least on a Megabase scale.^3,27–30^ Micro-C explores fine-scale structures of chromatin down to single nucleosome resolution.^31–33^ Genome folding studies via in-situ Hi-C across the tree of life in eukaryotes uncovered two types of genome architectures at chromosomal scale.^34^ Hi-C facilitated studies of chromatin organization reprogramming during early development stages after fertilization.^35^ Recently, PaleoHi-C revealed remarkable preservation of 3D genome architecture of a 52,000 year-old woolly mammoth (*Mammuthus primigenius*), proving the success of Hi-C on ancient, extinct samples.^36^

In Hi-C experiments, chromatin is first crosslinked with an agent like formaldehyde. Crosslinked DNA is then fragmented through enzymatic digestion. After labeling fragment ends with biotin, free fragment ends in physical proximity may ligate. After purification and shearing of DNA, ligated pair-ends are sequenced, and chromatin interaction data revealed by contacts populates the experiment’s outcome: the ligation map.

Despite successfully revealing genome architecture, many aspects of Hi-C experiments remain poorly understood. ^37–39^ Theoretical modeling, simulations and experiments have addressed some of these questions. For example, polymer collapse effects due to irreversible crosslinking between monomers on contact probabilities for different crosslinker concentrations were numerically modelled, and experimental contact probability curves for different crosslinking concentrations measured as a function of time in Hi-C experiments, studying polymer concentration evolution during crosslinking-induced collapse.^40^ Also, theoretical modeling of assay experiments were developed using chromatin polymer models to increase understanding on methods such as Hi-C.^41^ An interesting outstanding question concerns crosslinking, owing to different chemical efficiencies for direct crosslinking between DNA and DNA, DNA and protein, or protein and protein.^38^ One possibility suggests covalent bonding between spatially adjacent chromatin segments through protein bridges. If true, no clarity exists on the length of bridges binding pieces of DNA. Another possibility involves crosslinking DNA to an underlying protein matrix that percolates across the spatial extent of the nucleus, with long-range protein bridges serving as higher-order interactions between DNA and DNA.^38,42^ Evidence for this nuclear matrix as a polymer meshwork analogous to the cytoskeleton has been reported, although its function and existence remain an unsettled matter.^42–48^ Fig. 1 illustrates two limiting cases: crosslinking via short-range protein bridges (Fig. 1(A)), and the nuclear protein matrix (Fig. 1(B)).

**Figure 1.**
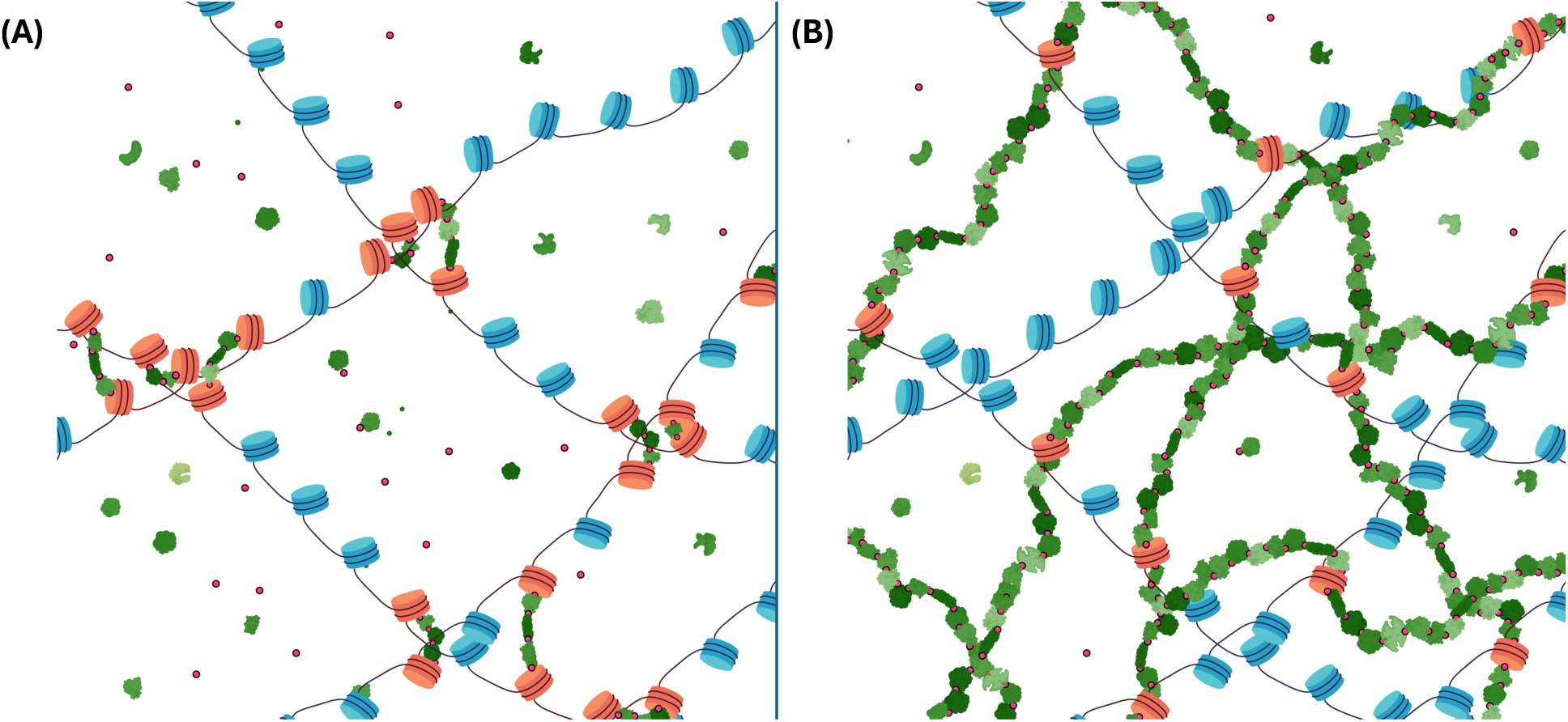
Two limiting cases of DNA crosslinking: Schematic representation of two limiting cases of crosslinking, in which DNA is represented by chains of nucleosomes (in blue), crosslinking agents such as formaldehyde are represented by small red dots and proteins are represented in different shades of green. Crosslinking can take place between the histones and proteins, as well as between proteins and other proteins. Crosslinked nucleosomes are represented in red. Two limiting cases are shown in (A) and (B), corresponding to crosslinking via short-range protein bridges, and crosslinking to a nuclear protein matrix, resp. ***(A) Short-range protein bridges:*** DNA is crosslinked to DNA via short-range protein bridges, allowing for a certain freedom of motion of the crosslinked nucleosomes in chromatin. Bridges shown include up to 3 proteins. ***(B) Nuclear protein matrix:*** a nuclear protein matrix is illustrated. DNA crosslinks to the scaffold provided by the protein polymer network that percolates the physical extent of the nucleus, analogous to the cytoskeleton in the cell. The protein meshwork can be understood as the long-range limit of the protein bridges, providing higher-order interactions that make up a network, providing some structural integrity and rigidity to the nucleus. Created in BioRender. Zubillaga, B. (2025) https://BioRender.com/aixb5aa

Open questions about the experiment’s chemical kinetics reveal uncertainty about effects of crosslinking agents, enzymatic digestion, and proximity ligation on Hi-C maps. Effects on maps due to effective rates of these reactions, non-equilibrium diffusive motion of digested fragments, and chromatin accessibility and density differences, require careful inquiry.

Ligation frequency dependence on genomic distance follows a characteristic power-law decay, reflecting complex long-range interactions in chromatin, illustrated by checkerboard patterns. However, one question not yet addressed is the dependence of ligation probability on 3D Euclidean distance, i.e., the ligation probability for a fragment end pair, originally separated by some Euclidean distance prior to digestion, as a function of said distance.

Maps are usually subjected to numerical post-processing and ad-hoc computational algorithms.^37^ Matrix balancing methods, such as Knight-Ruiz normalization, are ubiquitous when treating Hi-C maps.^49^ Closer inspection is warranted on their effects on raw data and our interpretation of post-processed maps.

Addressing some of these questions, we probe the inner workings of DNA-DNA proximity ligation assays through numerical experiments. We define an *in-silico* protocol representing Hi-C experiments, effectively modeling the sequence of chemical reactions of the actual experiment. This virtual Hi-C protocol mimics crosslinking of DNA to a percolating protein matrix, DNA binding via long-range protein bridges. DNA digestion with restriction enzymes or endonucleases and proximity ligation of free fragment ends are effectively modeled. Chromosome structures at nucleosome resolution are evolved in time with molecular dynamics and ligation events recorded populate ligation maps: the in-silico experiment’s output. In-silico maps agree with actual Hi-C experiments, showcasing features like plaid patterns, domains and compartments.

In-silico Hi-C grants control over basic chemical kinetic parameters, such as crosslinking, digestion, and ligation efficiencies, providing a window into their effects on ligation maps. The protocol can probe proximity ligation assays over ensembles of different structures (ensemble Hi-C) and single structures (scHi-C). It allows repetitions of in-silico experiments on a single structure, which cannot be accomplished on real, single cells in scHi-C.

We address the outstanding question of ligation probability with respect to initial Euclidean distances between loci pairs in the native conformations. Furthermore, we explore effects of numerical algorithms, such as Knight-Ruiz matrix balancing, on Hi-C maps from in-silico experiments, assessing post-processing effects on 3C methods.

## 2. IN-SILICO PROTOCOL FOR PROXIMITY LIGATION ASSAYS

We consider ensembles of structures 1.1 Mb in length, representative of chromosome 7 in human lymphoblastoid cells (genomic region: 95.4 to 96.5 Mbp), modeled at 200 bp resolution, for a total of 5500 beads per structure.^50^ Each bead represents a nucleosome and the DNA wrapped around it (accounting for ∼150 bp) plus linker DNA (∼50 bp). Prior to in-silico simulation of Hi-C, a preprocessing step on the ensemble relaxes each structure with the FIRE algorithm into energetically optimized states.^51^ The energy-optimized structures provide the ensemble of cells for in-silico Hi-C experiments.

Figure 2 illustrates the basic steps of in-silico Hi-C for an individual sample. A native structure is shown in Fig. 2.(A).(1) at nucleosome resolution, to be subjected to crosslinking, digestion and ligation. In Fig. 2.(A).(2), some nucleosomes (shown in red), picked at random according to a crosslinking efficiency, are crosslinked to the nuclear protein matrix. Some bonds, also picked at random according to an enzymatic digestion efficiency, are cut (depicted by straight orange lines). The protein matrix’s network-like nature confers a certain level of structural rigidity, implying, for the purposes of the in-silico protocol, that nucleosomes crosslinked to the matrix are rendered fixed in place inside the nucleus (their motions completely arrested). After crosslinking and digestion, resulting segment ends (shown in yellow and labelled A through H in Fig. 2.(A).(3)) can ligate under physical proximity. The crosslinked and digested structure is evolved in time with Langevin dynamics, according to a Hamiltonian including Lennard-Jones, FENE bonds and angle potentials. If, at some instant during time evolution, two segment ends are in physical proximity (within a certain threshold distance), a potential ligation event may happen according to a ligation rate (probability of ligation per unit time). If the ligation occurs, it is recorded as a contact and populates the ligation map.

**Figure 2.**
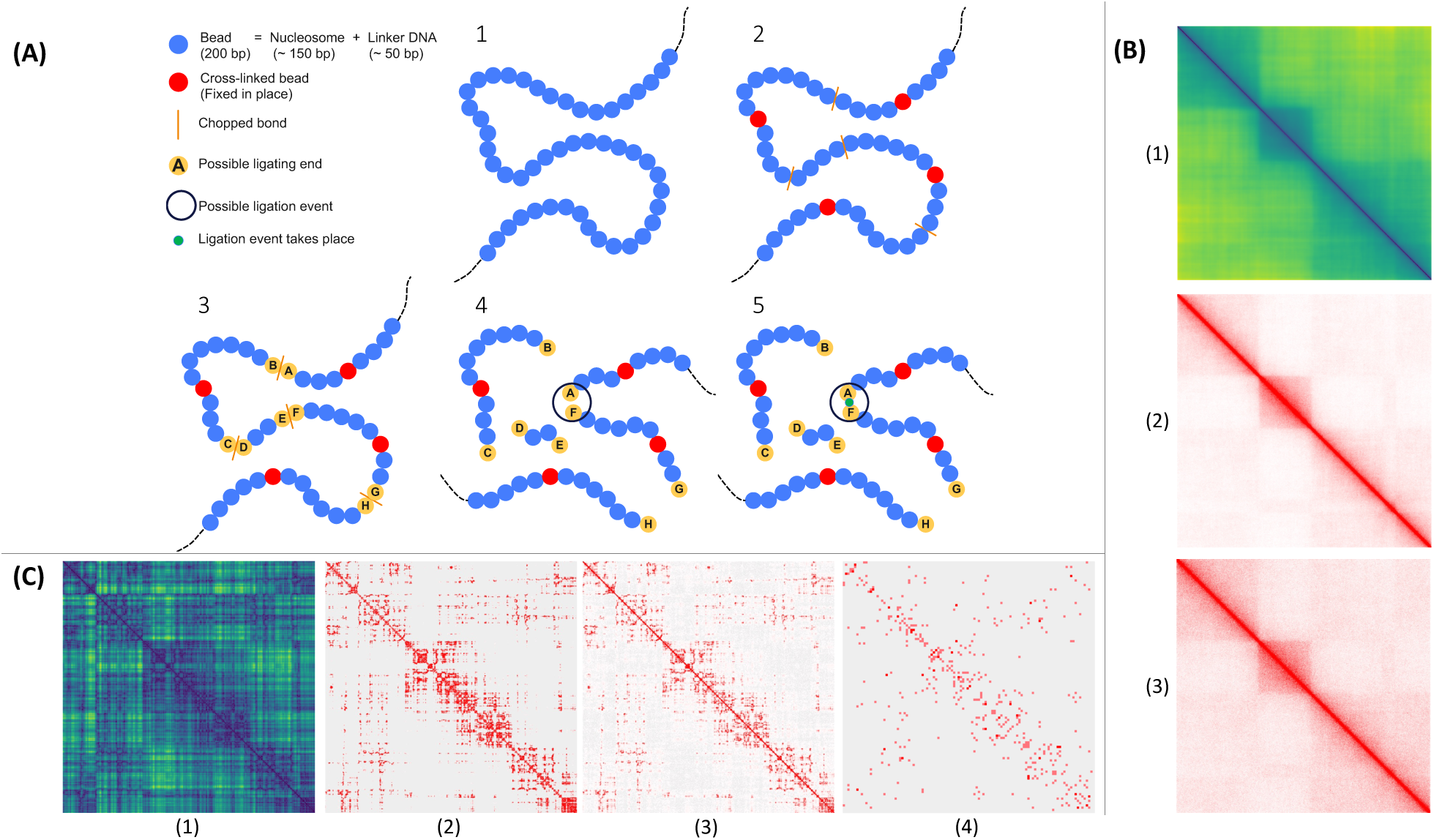
In-silico protocol at nucleosome resolution. ***(A) Basic steps:*** 1) Initial, native structure at nucleosome resolution (200bp). 2) Nucleosomes crosslink to protein matrix, fixed in place (in red). Bonds are enzymatically digested (cuts in orange). 3) Digested fragment ends (in yellow, labeled A through H) are free to ligate. 4) Structure evolves in time with molecular dynamics. 5) Ends A and F come into physical proximity (enclosed by circle). 6) Ends A and F ligate (green) with some probability rate. ***(B) Average distance, contact and ligation maps over native structure ensemble.*** Maps for ensemble of 5000 different structures modeling a 1.1Mbp region of chromosome 7 (95.4 to 96.5 Mbp) at 200bp resolution. (1) Distance map, in [nm], averaged over initial, native ensemble. (2) Average contact map of native ensemble. Beads pairs within distance of 𝑟 = 1.5𝜎 count as contacts. 𝜎 = 10 [𝑛𝑚] is the Lennard-Jones potential’s length-scale parameter. (3) Ligation map for 500 digested bonds, 500 crosslinked nucleosomes per structure, and ligation rate *p* = 10^−2^ *p*_0_, where *p*_0_ = 1/𝜏, and 𝜏 = 2.2678 ± 0.0008 [𝜇𝑠]. Proximity ligation is possible if two nucleosomes are within threshold distance of 𝑟 = 1.5𝜎. Correspondence between average distance and contact maps with ligation map shows protocol reproduces native ensemble features, including checkerboard patterns, domains and compartments. ***(C) Distance, contact and ligation maps of unique, single structure.*** Protocol is performed on individual structure of native ensemble. (1) Distance map, in [nm], of initial structure. (2) Contact map of initial structure. Contacts are counted for nucleosome pairs within threshold 𝑟 = 7.5𝜎, where 𝜎 = 10 [𝑛𝑚], as before. (3) Average ligation map over 5000 scHi-C iterations on same initial structure, with same numbers of digestions, crosslinks and ligation rate as before, and ligation threshold 𝑟 = 1.5𝜎. (4) Ligation map for single instance of in-silico scHi-C, at 10Kbp resolution for visual clarity, given the map’s sparsity at 200bp resolution (contrasted with Fig. 2.(C).(3)). Correspondence of distance and contact maps with ligation map is apparent. scHi-C protocol reproduces features of native structure and intimations of domains. In-silico protocol allows repetitions of experiment on same structure, aggregating over sparse single-iteration map, impossible in experimental scHi-C (single cells being single-use).

In-silico modeling of enzymatic digestion as a stochastic process, selecting bonds uniformly at random, cleaving them, and producing a randomly fragmented polymer, can be justified on the grounds of statistical analysis of human genomic sequences (see Supplementary Information Figs. S1 and S2). We searched the T2T reference genome for restriction sites corresponding to 4-cutter enzyme MboI and to 6-cutter enzymes HindIII and NcoI. We found the sites to be uniformly distributed across the genome and distances between successive restriction sites to be exponentially distributed, consistent with uniform digestion.

This in-silico protocol enables calculations of ligation maps and ligations frequencies as functions of both genomic and Euclidean distances. It allows effective characterizations of the basic chemical kinetics of the experiment, i.e., effects of crosslinking, enzymatic digestion, and ligation efficiencies on ligation maps. It also enables assessments of numerical post-processing algorithms (such as Knight-Ruiz matrix balancing) on ligation maps in contrast with raw, unprocessed data.

## 3. RESULTS

### In-silico proximity ligation assays, crosslinking to a percolating protein matrix, successfully replicate ensemble and single-cell Hi-C experiments

In Fig. 2, we present results from the virtual Hi-C protocol on the ensemble of native structures at a nucleosome resolution. Fig. 2.(B).(1) shows the average distance map of the ensemble. The ensemble of native structures successfully reproduces main characteristic of real Hi-C maps, such as high numbers of counts along the diagonal (due to local interactions between nucleosomes separated by short genomic distances), checkerboard patterns, domains and compartments. Said features are also present in the average contact map on the ensemble of native structures (Fig. 2.(B).(2)), in which a contact is only counted if the Euclidean distance between its corresponding pair of loci is below a certain threshold. Fig. 2.(B).(3) shows the final ligation map of the in-silico Hi-C experiment (after simulating crosslinking, digestion and ligation) averaged over all structures in the ensemble. Compared with the average distance and contact maps, the ligation map closely resembles the native ensemble of structures, successfully representing Hi-C in-silico.

In-silico Hi-C can also simulate scHi-C methods, enacting the protocol over a single, initial native structure, as shown in Fig. 2.(C). An interesting possibility of this numerical protocol, in contrast with actual scHi-C experiments, is that it can be carried out multiple times over the same initial structure and the results aggregated to produce a higher total number of contacts and a richer ligation map. This, of course, is a luxury that cannot be afforded in actual scHi-C experiments, where a single cell can only be probed once, severely limiting the contact yield of the final scHi-C maps. Fig. 2.(C).(1) shows the distance map of the single, initial structure, where intimations of domains and compartments in the sample can be appreciated. These features are appreciable in Fig. 2.(C).(2), which shows the contact map for the single structure, consistent with the distance map. As expected for a single structure, the contact map shows considerable sparsity compared to its ensemble counterpart in Fig. 2.(B).(2).

Fig. 2.(B).(3) shows the ligation map resulting from repeating the protocol over the single initial structure many times and aggregating the results from each iteration. The sampling of crosslinked loci and digested bonds, as well as the physical trajectories and ligated ends that ensue through time evolution, are independent across all repetitions of the experiment on the single native structure. There are notable differences between aggregating multiple iterations on the same initial structure, Fig. 2.(B).(3), and simulating the protocol over said structure just once, as shown in Fig 2.(B).(4), where sparsity is notorious. Repeating the protocol multiple times over the same native structure and aggregating the results makes the features of scHi-C maps clearer. Differences between the ensemble Hi-C and the scHi-C protocol, as contrasted by Figs. 2.(B) and 2.(C), are consistent with real experimental ligation maps, where sparsity of the latter with respect to the former is systematic, owing to the considerable variability between chromosome conformations amongst individual cells in the ensemble.

Further evidence of consistency with experiment is provided by the probability of ligation as a function of genomic separation between pairs of loci in Fig. 3.(A). It is well-established by experiment that contact probabilities follow characteristic power-law scalings, reflecting the complexity of long-range interactions (domains, loops and compartments). Fig. 3.(A) shows the ligation frequency versus genomic distance for the native ensemble, whereas Fig. 3.(B) does so for a single structure sampled many times (same structure as in Fig. 2.(C)). In both cases, the ligation frequency curves are ordered according to the ligation rate, varied across orders of magnitude. The in-silico protocol recovers scaling behavior of the ligation frequency, consistent with experiment. Fig. 3.(A) shows ligation frequencies versus genomic distance for different ligation rates (probability of ligation per unit time), agreeing with the underlying contact probability scaling of the ensemble of initial native structures. Fig. 3.(E) shows excellent agreement with actual experimental ligation frequencies from a Hi-C map at 1Kbp resolution, corresponding to the same region of chromosome 7 (genomic region: 95.4 to 96.5 Mbp).

**Figure 3.**
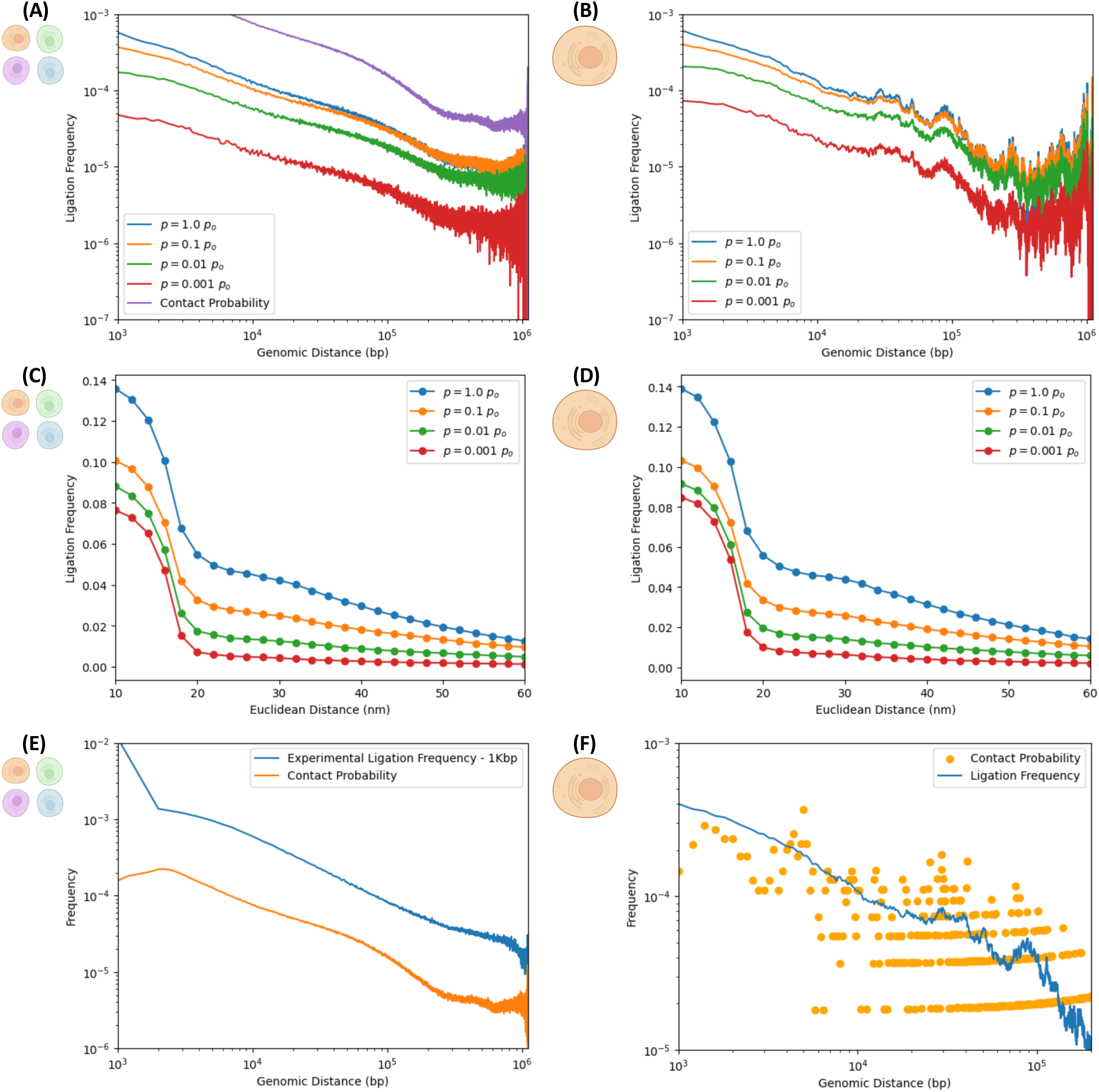
Dependence of ligation frequencies on genomic and Euclidean distances. Left and right columns correspond to numerical experiments simulating ensemble and single-cell Hi-C experiments, resp. ***Ligation frequency as a function of genomic distance for different ligation rates:*** Ligation frequencies exhibit characteristic power-law scaling (typical of experimental maps) for different ligation rates “*p”,* capturing long-range interaction effects of features as domains and compartments. Results shown for different ligation rates span a few orders of magnitude both for: (A) ensemble Hi-C calculations (on 5000 different structures) and (B) scHi-C calculations (aggregating 5000 realizations on a single structure). For comparison, the average contact probability of the ensemble is shown.^50^ In (A), power-law scalings for different ligation rates resemble the contact probability over the native ensemble of initial structures. ***Ligation frequency as a function of the 3D Euclidean distance between pairs of nucleosomes for different ligation rates:*** Ligation frequency versus the Euclidean distance between pairs of loci exhibits characteristic sigmoidal decay, suggesting a characteristic length scale, in the order of ∼20 [nm], over which most of the ligation events occur. Ligation frequency at Euclidean distance *“d”* represents the frequency with which pairs of free segment ends, initially separated by Euclidean distance *“d”* in the initial, native conformations, find themselves within physical proximity (within a distance of 𝑟 = 1.5𝜎) at some point in time and ligate according to the ligation rate. These sigmoid curves are not proper probability distribution functions and not subject to normalization. Results shown for different ligation rates span a few orders of magnitude both for: (C) ensemble Hi-C (on 5000 different structures) and (D) scHi-C (repeated 5000 times on same structure). ***Agreement with experimental contact probability:*** (E) The ensemble contact probability over 5000 native structures representative of a 1.1Mbp region of chromosome 7 agrees with corresponding experimental results over same region at 1Kbp resolution, sharing similar exponents. ***Averaging effect in scHi-C simulations:*** (F) Average ligation frequency from 5000 iterations of single-cell numerical experiment on the same initial structure, contrasted with underlying contact probability of said structure. Averaging on different realizations reduces ligation frequencies noisiness relative to the initial structure’s contact probability. Created in BioRender. Zubillaga, B. (2025) https://BioRender.com/6afacus

As expected, the ensemble calculation shows smoother power-law scaling than their single-cell counterparts. This is a consequence of the large variability between different samples (cells) and the effect of averaging over ensembles of different cells with sample-to-sample fluctuations, apparent by comparison between ensemble and single-cell ligation maps. Indeed, Figure 3.(B) still presents evidence of power-law scaling, albeit with greater noise on account of the sparsity of the scHi-C map.

The noisy curves in Fig. 3(B) already manifest a smoothing effect of their own, since they aggregate results of many iterations of in-silico scHi-C experiments on the same initial, native structure. The sets of nucleosomes and bonds selected for crosslinking and cleavage, resp., in each realization are statistically independent from those in other iterations. Subdiffusive trajectories that ensue from time evolution with Langevin dynamics are different from one iteration to another. This provides variability on ligation events that get recorded in different iterations, although they all sample the same initial structure. Hence, variability provides some measure of smoothness after results from the different iterations are aggregated. Figure 3.(F) illustrates the smoothing effect by comparing the single-cell ligation frequency averaged over all iterations with the underlying contact probability of the initial, native structure. The latter is much noisier than the former, illustrating the role that non-equilibrium effects of distinct subdiffusive trajectories can have in averaging out iteration-to-iteration fluctuations, even if all iterations probe the same native sample.

### Crosslinking chromatin to underlying protein network produces maps consistent with experimental results

Results discussed thus far in Figs. 2 and 3 show that the in-silico protocol, assuming the existence of a nuclear matrix, is consistent with experimental results from actual 3C experiments. Ligation maps with checkerboard patterns, domains and compartments, are consistent with the average contact and Euclidean maps of the ensemble of native structures; and ligation probabilities versus genomic distance recover the expected power-law scaling. This consistency, assuming the protein meshwork as a backdrop, does not rule out other possibilities on the issue of crosslinking, as the opposite limiting case of short-range protein bridges could also produce similarly consistent results.

### In-silico Hi-C protocol enables estimation of ligation frequency as a function of Euclidean distance, revealing the existence of a typical length scale for ligation events

Although Hi-C experiments can estimate the ligation frequency as a function of genomic distance, a different but related question remains unanswered: what is the ligation frequency as a function of Euclidean distance? In Figs. 3.(C) and 3.(D), we address this question with the in-silico protocol. A data point in one of these curves represents the probability that two free segment ends, initially separated by a Euclidean distance *“d”* in the native structure, will, at some point in time, find themselves in physical proximity (within a distance of 𝑟 = 1.5𝜎 = 15 [nm]) and ligate with a ligation rate *“p”* (probability of ligation per unit time). Ligation frequency as a function of Euclidean distance exhibits a sigmoidal shape, which suggests a characteristic length scale within which the ligation of free segment ends take place, typically of the order of some 20 [nm] or so. This is consistent with the proximity-based character of 3C methods, in which digested segment ends in physical proximity may bind through a ligation reaction, favoring ligations between loci that are close together in Euclidean space in the native conformations versus ligations between pairs of loci that are distant. Fragments ends corresponding to two distant loci in 3D would have to come into close physical proximity at some point in the sub-diffusive time evolution, which is less likely than encounters between initially proximate fragment ends in the native structures. The calculation of the sigmoid, revealing the local nature of proximity ligation events and estimating the characteristic length scale for ligation events, are results that have not yet been addressed experimentally, made possible by harnessing the power of in-silico experiments.

### Non-equilibrium effects manifest through changes in power-law scaling exponents of ligation frequencies versus genomic distance

To assess the effect of basic chemical kinetics parameters on the results of proximity ligation assays such as ensemble Hi-C, we perform in-silico experiments for different digestion efficiencies and ligation rates, fixing the number of crosslinked nucleosomes. Figs 4.(A) and 4.(B) illustrate the power-law scaling of the ligation frequencies versus genomic distance for the fastest and slowest ligation rates, resp., explored in this work, i.e., *p* = 1 × *p*_0_ and *p* = 10^−3^ × *p*_0_. In each case, plots are shown for different digestion efficiencies, measured by the total number of bonds cleaved or cut (ranging from 500 to 5000 digestions per sample), mimicking the effect of restriction enzymes or endonucleases. For “fast” ligation efficiencies (Fig. 4.(A)), the curves for different digestion efficiencies reveal very similar behaviors and scaling exponents, similar to the contact probability of the underlying ensemble of native structures. However, for “slow” ligation efficiencies (Fig. 4.(B)) the scaling exponent changes appreciably with the number of bonds digested. Increasing number of digested bonds, the power laws progressively flatten out and the exponent decreases. The apparent insensitivity to the number of digestions for the “fast” ligation rate is a consequence of ligation events taking place very quickly, individual ligated fragments prevented from moving or diffusing away from their initial positions inside the nucleus significantly. Hence efficiently captured ligations occur between fragment ends that resemble the chromosome conformations of the native structures. For the “slow” ligation rates, a potential ligation event is a thousand times less likely to materialize, thus providing time for fragments to move and diffuse away from their initial positions in the native structures. This non-equilibrium effect of subdiffusive motion becomes more apparent with increasing numbers of bonds digested, thus changing the exponent of the power-law scalings as curves flatten with greater numbers of digested bonds.

**Figure 4.**
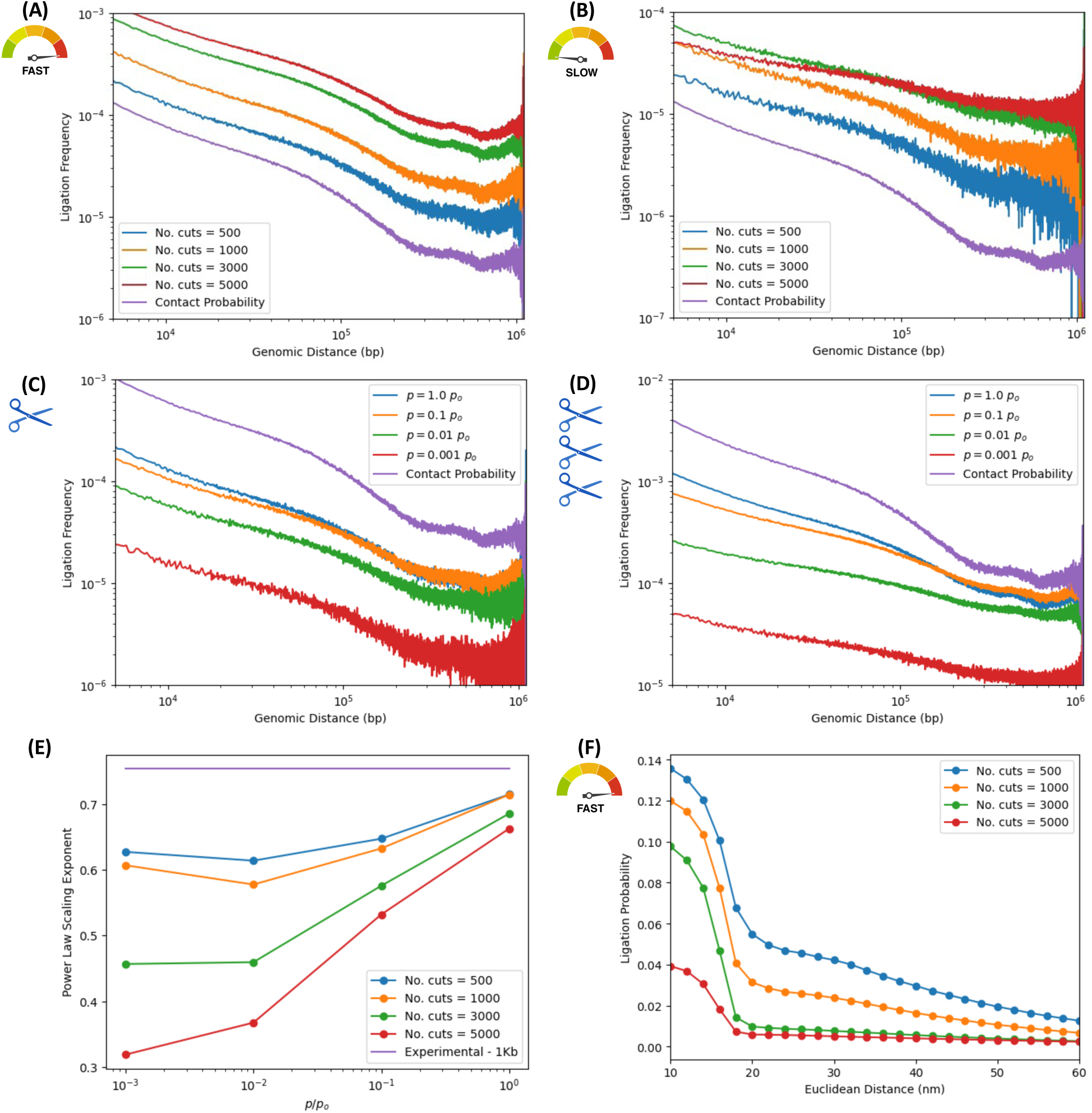
Digestion efficiency effects on ligation frequencies for different ligation rates. In-silico ensemble Hi-C for different digestion and ligation efficiencies and 500 crosslinks. “No. cuts” is the number of bonds enzymatically cleaved, spanning the range from 500 to 5000. Ligation rates span orders of magnitude. Contact probability of native ensemble is shown for comparison. ***Ligation frequency vs. genomic distance for different digestion efficiencies:*** We explore non-equilibrium effects of fragment diffusion on ligation maps, in fast and slow ligation rate limits, for different digestion efficiencies. (A) Fast ligation rate (*p* = 1 × *p*_0_): Similar power-law exponents, irrespective of digestion efficiency. Ligations accumulate quickly, individual fragments not diffusing significantly from initial positions before ligating. Hence the insensitivity to digestion efficiency. (B) Slow ligation rate (*p* = 10^−3^ × *p*_0_): Exponents change appreciably, and curves “flatten”. Greater fragmentation (i.e., smaller fragment sizes) progressively degrades ligation map due to non-equilibrium effects of fragment diffusion. ***Ligation frequency vs. genomic distance for different ligation rates:*** Ligation frequencies versus genomic distance are shown for high and low digestion efficiency and different ligation rates. (C) Low digestion efficiency: Just 500 bonds digested, i.e., 9.09% of bonds cleaved, for average fragment sizes of ∼11 nucleosomes. Exponents are rather insensitive to ligation rates. Low cleaving rate implies low structural degradation for the ligation map. (D) High digestion efficiency: 5000 bonds digested, i.e., 90.9% of bonds cleaved, for average fragment sizes of ∼1.1 nucleosomes, near nucleosome gas limit. The polymer is almost fragmented into individual nucleosomes that diffuse subject to excluded volume. After degrading chromosomes into nucleosome gases, only large ligation rates preserve information in maps. For low ligation rates, gas diffusion effaces structure. Scaling laws flatten with decreasing ligation rates for highly digested structures. ***Effect of digestion efficiencies on power-law exponents of ligation frequencies vs. genomic distance:*** (E) Non-equilibrium effects of digestion efficiency on exponents for different ligation rates. Curves for different numbers of bonds cleaved are shown. Exponents decrease with decreasing ligation efficiency, as suggested in (A) through (D). (F) Sigmoidal decay of ligation frequency versus Euclidean suggests a characteristic distance of ∼20 [nm] when *p* = 1 × *p*_0_. Ligation frequencies are ordered according to number of cuts, with large digestion efficiency corresponding to lower frequencies. Created in BioRender. Zubillaga, B. (2025) https://BioRender.com/pc4u4ij

This non-equilibrium effect is also shown in Figs 4.(C) and 4.(D), where ligation frequencies versus genomic distances are shown for the lowest and the highest digestion efficiencies explored in the work, resp., i.e, 500 and 5000 bond cuts.

Fig. 4.(C), corresponding to the lowest digestion efficiency, shows the ligation frequency scaling laws for several ligation rates (spanning orders of magnitude). Since only 500 digestions take place for a given sample, corresponding to ∼9% of linker DNA cleaved, average fragment sizes are some ∼11 nucleosomes in length, and these fragment lengths (relative to the fixed number of 500 crosslinks) can preserve the structure of the conformations of the initial native ensemble.

Fig. 4.(D), however, corresponds to the highest digestion efficiency (for the same set of ligation rates). Given that 5000 bonds are now digested, corresponding to ∼90% of the linker DNA cleaved, average fragment sizes are now ∼1 nucleosome in length, i.e., a “gas” of nucleosomes obtains, which diffuses away (with dynamics approaching normal diffusion), effacing structure from the Hi-C map as nucleosomes move away from their initial positions in the native structures. Greater fragmentation of the polymer corresponding to smaller average fragments, without a corresponding increase in the number of crosslinked beads, leads to increasing degradation of the chromatin structure captured by ligation maps and the ligation frequencies. This progressive effacement of structure pushes the Hi-C map towards a constant matrix with increasing degradation; hence the flattening of the power-law scalings.

To better assess the non-equilibrium effects of the subdiffusive motion, Fig. 4.(E) quantifies the power-law exponent of ligation frequencies versus genomic distance. These exponents are obtained from least-squares fits to power laws for range of ligation rates (spanning orders of magnitude) and digestion efficiencies (from ∼9% to ∼90% of linker DNA cleaved), maintain the same fixed crosslinking efficiency throughout. Curves corresponding to different digestion efficiencies, show the exponent as a function of ligation rate. As before, fast ligation rates permit a quick accumulation of ligation events and an efficient population of the ligation map, thereby reflecting the structure of the original chromosome conformations. Slow ligation rates imply a greater time scale for ligation events to occur, thereby allowing diffusive effects to efface structure as fragments displace away from their original positions. As before, the degradation effect becomes greater with increasing digestion efficiency, since average fragment become smaller, and the diffusion of small fragments effaces information of the 3D architecture of the native ensemble. The greatest degradation of the power-law exponent is consistent, therefore, with the lowest ligation rate, in the nucleosome gas limit. Further exploration of non-equilibrium effects of subdiffusive motion of DNA fragments and their degrading effects on the power-law scalings of the ligation frequencies is presented in the Supplementary Information (Figs. S3, S4 and S5).

Numerical estimations of degradation effects in ancient DNA samples on contact probability scalings due to diffusive dynamics on a nucleosome “gas” were studied to understand the remarkable preservation of genome architecture in PaleoHi-C experiments on woolly mammoth skin cells.^36^ Results on diffusion models of degradation on nucleosome gases are consistent with the findings shown in Fig. 4.

### Increasing concentration of crosslinks to nuclear protein matrix helps preserve 3D genome architecture

Fig. 5 illustrates effects of varying crosslinking agent concentrations at the beginning of the experiment. Figs. 5.(A) and 5.(B) present ligation frequencies versus genomic distance for several crosslinking efficiencies in the fast and small ligation rate limits, resp., for fixed bond digestion efficiency. In each case, the crosslinking efficiency range explored goes from the limit no crosslinking whatsoever to the opposite extreme where all nucleosomes are fixed in place, crosslinked to the protein network.

**Figure 5.**
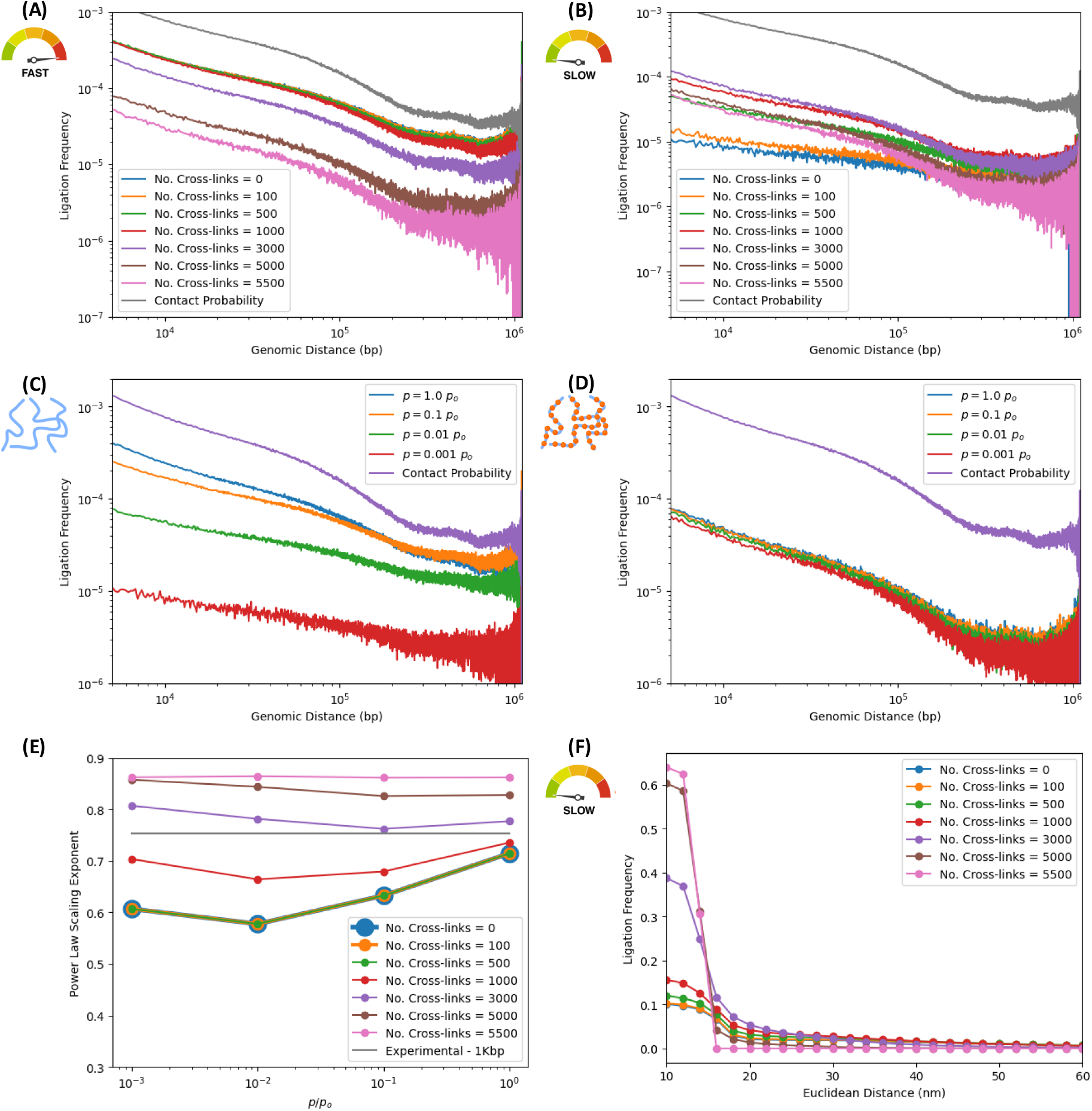
Crosslinking efficiency effects for different ligation rates. Ensemble protocol for different crosslinking and ligation efficiencies, and 1000 digested bonds. “No. Cross-links” represents the number of crosslinked nucleosomes, spanning the range from 0 (no crosslinking) to 5500 (fully crosslinked, frozen structures). Ligation rates span a few orders of magnitude. For comparison, the contact probability of native ensemble is shown. ***Ligation frequency vs. genomic distance for different ligation rates:*** We explore fast and slow ligation rate limits, for different crosslinking efficiencies. (A) Fast ligation rate *p* = 1 × *p*_0_: Scaling laws have similar exponents irrespective of crosslinking efficiency. For high ligation rates, ligations accumulate on short timescales, fragments not diffusing significantly before ligating. (B) Slow ligation rate *p* = 10^−3^ × *p*_0_: Power-laws “flatten” with decreasing crosslinking efficiency. Small ligation rates imply slower ligation accumulation and non-equilibrium effects of fragment diffusion impacting the map. ***Ligation vs. genomic distance for different crosslinking efficiencies:*** We consider low and high crosslinking efficiencies, for different ligation rates. In the former, no nucleosomes are crosslinked, and fragments can diffuse away, subject to excluded volume. In the latter, 5000 nucleosomes (i.e., 90.9% of the nucleosomes) are crosslinked to the matrix. (C) No crosslinking: In the absence of crosslinks, power-laws flatten with decreasing ligation efficiencies as structure washes away from the map with increasing and unimpeded fragment diffusion. (D) High crosslinking limit: Structures are frozen in their initial, native states. Crosslinks arresting fragment motion makes ligation frequencies insensitive to ligation rates and map impervious to diffusion. ***Digestion efficiency effects on power-law exponents of ligation frequencies vs. genomic distance:*** The effects just described are quantitatively measured by the scaling exponent vs. ligation rate for different crosslinking efficiencies (E) For low crosslinking efficiencies (0 to 500 crosslinks, i.e. 0% to 9.09% of nucleosomes crosslinked), the exponent decays with decreasing ligation efficiencies. For high crosslinking efficiencies, structures are effectively frozen in place and the exponent is insensitive to ligation rates. ***Effect of crosslinking efficiency on the ligation frequency vs. Euclidean distance:*** (F) Ligation frequency decays sigmoidally with a typical distance of ∼15[nm] for different crosslinking efficiencies and a low ligation rate *p* = 10^−3^ × *p*_0_. Ligation frequencies are ordered according to crosslinking efficiency. For lower crosslinking, unhampered fragment motion permits ligations between initially distant fragments. In fully crosslinked structures, fragment motion is arrested ligations between distant fragments being exceedingly unlikely, causing sharp sigmoidal decays. Created in BioRender. Zubillaga, B. (2025) https://BioRender.com/eir7kx6

Fig. 5.(A) shows that, in the fast ligation rate limit (*p* = 1 × *p*_0_), the power-law exponent remains largely insensitive to crosslinking concentration changes (bearing excellent agreement with native structure ensemble’s contact probability), as the typical time scale for ligations to occur is small and they accumulate early on while fragments haven’t yet diffused significantly.

However, for slow ligation efficiencies (*p* = 10^−3^ × *p*_0_), the role of crosslinks between fragments and the protein matrix become evident (Fig. 5.(B)). Greater crosslinking efficiencies, measured by the number of nucleosomes crosslinked, lead to better structural preservation of initial conformations (fragment motions effectively arrested with high crosslinker concentrations), as seen by comparing different power-law scalings curves with the native ensemble’s contact probability. Smaller crosslinking concentrations allow for greater numbers of digested fragments disconnected from the protein meshwork and displacing away from their initial positions in the native conformations. This non-equilibrium effect due to subdiffusive dynamics of free fragments leads to structural information loss in ligation maps, and produces a flattening effect in the power-law curves (compared to the contact probability of the native ensemble). Lowering crosslinker concentrations (relative to fixed ligation and digestion efficiencies) leads to progressive deterioration ligation map quality.

This is also appreciated in Figs. 5.(C) and 5.(D), where ligation frequencies are presented, for different ligation rates (spanning orders of magnitude), resp., in the limits of no crosslinking and of high crosslinker concentration (where ∼91% of nucleosomes in the sample crosslinked to the matrix). In the absence of crosslinking, information degradation manifests as a flattening of power laws with increasingly slower ligation rates (relative to the contact probability of the native ensemble.) In the opposite limit of high crosslinking efficiencies, most nucleosomes are pinned down to the protein meshwork and fragment motion is effectively arrested, leading to remarkable preservation of structure, with excellent agreement between power-law exponents of ligation and contact probabilities.

Fig. 5.(E) quantifies the effect crosslinking efficiency on exponents for different ligation rates, with fixed digestion efficiencies. For low crosslinker concentrations (less than 10% of the nucleosomes crosslinked to the matrix), exponents are insensitive to the crosslinking agent. In this case, the fragments’ motions are effectively unimpeded by the protein meshwork, and smaller exponents in the power-law scaling are observed.

In the limit of high crosslinker concentration, exponents become well preserved due to high numbers of fragments pinned down to the protein network, conserving information about the three-dimensional conformations of native samples.

This high crosslinking limit yields ligation frequencies versus Euclidean distance that show the typical sigmoidal shape discussed previously (Supplementary Information Fig. S6), with very high frequencies for small distances decaying sharply beyond the characteristic length scale. Due to the arrested motion of nucleosomes, not displacing significantly from their initial positions in the native conformations, information is preserved. The effect of varying the crosslinking efficiency on the sigmoidal decay, in the slow ligation rate limit for a fixed digestion efficiency, is shown in Fig. 5.(F). The arresting effect of high crosslinking concentrations shows sharply decaying sigmoids (fully crosslinked DNA) progressively softening into smoothly decaying curves as crosslinking efficiency is lowered and fragments motions are permitted. A broadening of the curves ensues, as diffusive motion permitting ligations between fragment ends that have displaced away from their original positions in the native conformations.

Further exploration on the role of non-equilibrium effects of fragment diffusion and the motion-arresting effects of crosslinks to the nuclear protein matrix on contact maps and contact probabilities and power-law scaling is presented in the Supplementary Information (Figs. 7, 8 and 9).

### Effect of numerical postprocessing of ligation maps through Knight-Ruiz balancing algorithm is minor relative to raw data

A popular practice in 3C, numerical post-processing algorithms balance Hi-C maps to enforce double-stochasticity, all individual rows and columns adding to the same number of ligations.^37,49,52^ A mean-field assumption of sorts, individual loci are thought to have similar average numbers of contacts. If said average is statistically significant, locus-to-locus fluctuations in ligation numbers per locus are negligible leading to statistical homogeneity across the chromosome. Exploring numerical post-processing effects, we compare results from raw in-silico maps with their post-processed counterparts, balanced with the Knight-Ruiz (KR) algorithm (a computationally efficient method for symmetric matrix normalization).^49^ Fig. 6 contrasts raw data from non-normalized maps with their KR-normalized counterparts.

**Figure 6.**
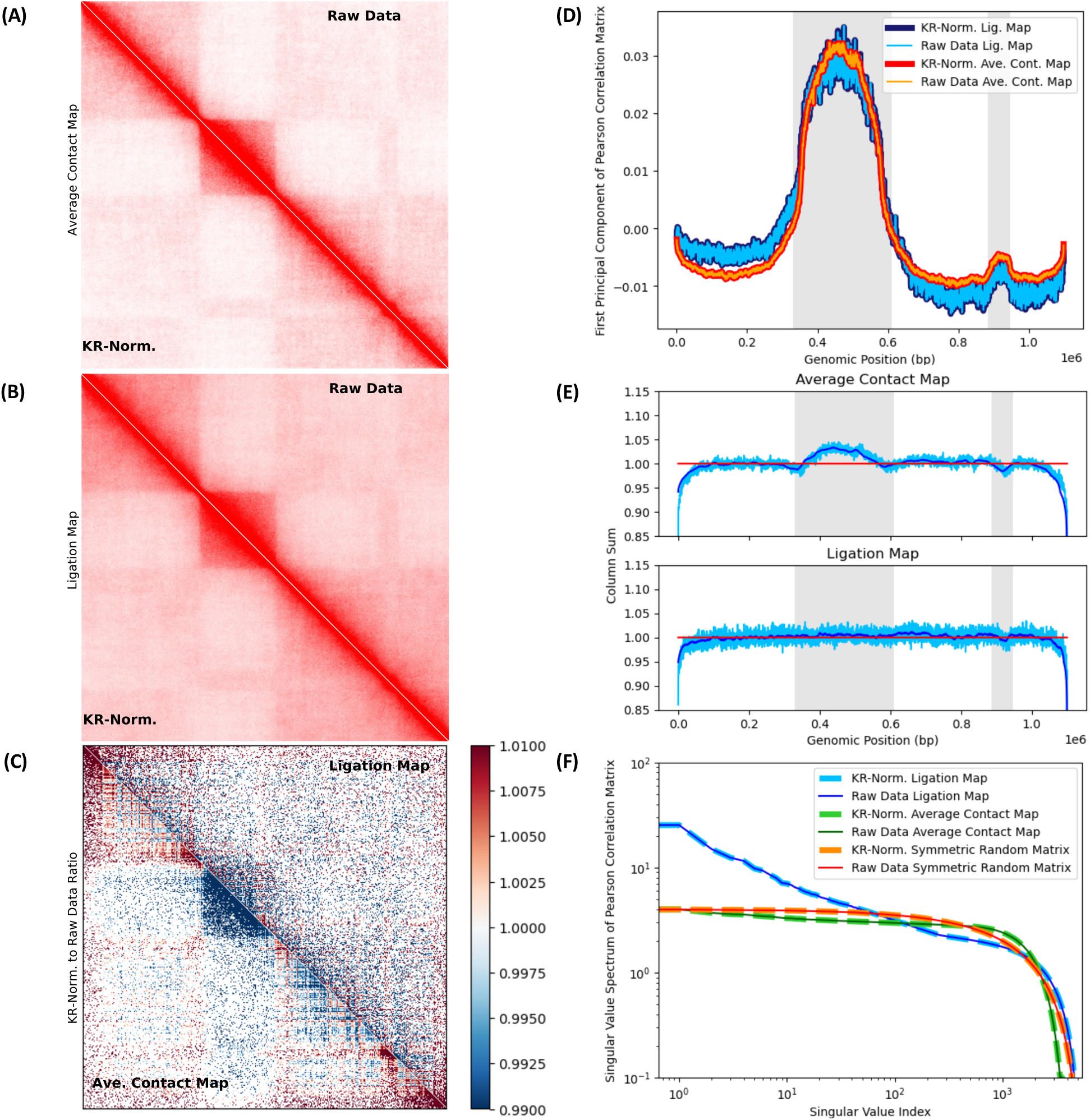
Knight-Ruiz (KR) normalization effects on Hi-C maps. Matrix balancing effects are studied, contrasting KR-normalized (post-processed) with non-normalized (raw) maps. In-silico ligation rate *p* = 10^−1^ × *p*_0_, 3000 bond cuts and 500 crosslinks are considered. ***(A) KR-normalized and raw average contact maps.*** The upper triangular matrix (UTM) shows the raw contact map on the native ensemble. The lower triangular matrix (LTM) shows its KR-normalized counterpart. No visually obvious difference exists between them. ***(B) KR-normalized and raw ligation maps.*** The UTM and LTM show raw and KR-normalized ligation maps, resp. No visual difference is apparent. ***(C) KR-normalized to raw matrix ratio for ligation and average contact maps.*** The UTM and LTM show the ratio of KR-normalized to non-normalized matrices for ligation and contact maps, resp. Normalization depletes the contact map’s central domain (dark blue), enriching leftmost and rightmost domains (red). The corresponding effect on the ligation map is less striking, because of non-equilibrium effects of diffusion after fragmentation, free segments displacing before ligating. ***(D) First Principal Component (FPC) of Pearson Correlation Matrix (PCM).*** The FPC of the PCM is shown for raw and KR-normalized maps, for both average contact and ligation maps. Shaded backgrounds (gray) indicate compartment switches. In both ligation and contact maps, FPC captures compartment switches (see 6.(A) and 6.(B)). There is no significant difference between FPCs of raw and KR-normalized data, as their curves overlap. ***(E) Column sums for average contact and ligation maps.*** Column sum per locus is shown versus genomic position for contact and ligation maps (above and below, resp.) Pre and post normalization sums (light blue and red, resp.), and moving averages over the former (dark blue) are shown. Normalization succeeds, column sums converging to 1. Column sums over contact map distinguish compartment switches (see gray shaded backgrounds). A visible enrichment around the central domain, as well as a small depletion, agree with the FPC in 6.(D). Noisier column sums for ligation map lack obvious enrichments because of non-equilibrium effects. ***(F) Singular value spectra of PCMs.*** Spectra for KR-normalized and non-normalized data, for ligation and contact maps, and a random symmetric matrix with entries drawn from *U*(0,1), show insensitivity to normalization.

In Figs. 6.(A) and 6.(B), upper triangular matrices show maps corresponding to raw data, while lower triangular matrices show their KR-normalized counterparts. Figs. 6.(A) and 6.(B) show, resp., effects of matrix balancing on average contact maps (over the native structure ensemble) and on in-silico Hi-C ligation maps (after crosslinking, digestion, time evolution and ligation). Visual examinations of raw versus corresponding KR-balanced maps do not reveal obvious differences, suggesting that perhaps numerical normalization is justified.

The matrix in Fig. 6.(C) shows the ratio of KR-normalized maps to their corresponding raw data maps. The upper triangular matrix shows the ratio for the ligation map, while the lower triangular matrix shows the same ratio for the average contact map on the native structure ensemble. On the average contact map (not subjected to in-silico Hi-C), KR-balancing drains the contact numbers in the central domain, accompanied by a corresponding increase in the leftmost and rightmost domains. This effect is less evident in the ligation map shown, due to non-equilibrium effects from digestion and diffusion of free fragments.

In Fig. 6, we perform Principal Component Analysis (PCA) on the Pearson Correlation Matrix (PCM) of ligation and average contact maps, both before and after KR-normalization. Fig. 6.(D) shows the First Principal Component (FPC) for raw and KR-normalized maps. The FPCs of ligation and average contact maps successfully capture compartment switches (Figs. 6.(A) and 6.(B)). There is no considerable difference between the FPCs of raw and KR-normalized maps, since curves of raw and balanced data overlap, both for ligation and average contact maps. As far as FPCs of PCMs are concerned, KR-normalization does not significantly distort the information in the raw maps.

In Fig. 6.(E), we show the map’s column sums for each locus versus genomic position, both for the ligation and average contact maps. The column sum (light blue) shows the magnitude of locus-to-locus fluctuations relative to the moving average (dark blue). The KR algorithm successfully normalizes the matrices and balanced column sums converge to 1 (shown in red). Column sums for the average contact map, prior to normalization, distinguish compartment switches (alternating white and gray shades). The central domain’s visible enrichment and the small depletion to the right are consistent with the pattern revealed by the FPC in Fig. 6.(D). KR-balancing also normalizes the ligation map, but raw data is noisier because of non-equilibrium effects of diffusive motion, free segments displacing from their original positions in the undigested chromosome until ligations take place. Averages contact maps on crosslinked and digested structures, interrogated at different stages of diffusion, reveal non-equilibrium effects of fragment motions on progressive effacement of enriched column sums in the central domain (see Supplementary Information S10).

Singular Value Decomposition (SVD) of the PCM is performed, before and after KR-balancing of the maps, and singular value spectra are shown for ligation and average contact maps in Fig. 6.(F). For comparison, we show results for a random symmetric matrix with entries drawn from a standard uniform distribution *U*(0,1). In all cases, singular value spectra remain unaffected after KR-normalization.

Insensitivity of FPCs and SVD spectra to KR matrix-balancing and striking visual similarity between pre and post-processed maps, suggest consistency with the mean-field assumption. However, although there is no apparent difference between the pre and post-normalized maps, other metrics, such as ratios of KR-normalized to raw maps and column sums of the maps, are affected: count-enriched domains depleted at the expense of enrichment in other parts of the map.

## 4. DISCUSSION AND CONCLUSIONS

We developed an in-silico protocol simulating proximity ligation assays for 3C with molecular dynamics and physical modeling, mimicking effects of DNA crosslinking, digestion and fragmentation of crosslinked DNA with restrictions enzymes or endonucleases, and subsequent ligations between fragment-end pairs. The protocol allows control over basic variables affecting the chemical kinetics of Hi-C experiments, such as crosslinking, digestion and ligation efficiencies, thus aiding our understanding on inner workings and gray areas of the assay. Simulated on ensembles of native structures representative of human lymphoblastoid cells, it successfully reproduces typical features of Hi-C maps, such as checkerboard patterns, domains and compartments, replicating characteristic power-law decay of ligation frequencies versus genomic distance.

In addition to ensemble Hi-C, the protocol also simulates scHi-C methods, with the added luxury of enabling multiple repetitions of the single-cell experiment on the same initial structure, aggregating over all iterations. It can assess the role of averaging, both over multiple iterations on a single sample and on realizations on ensembles of different structures, in softening noisiness of ligation maps and frequencies.

Non-equilibrium effects due to subdiffusive time evolution of crosslinked and digested structures manifest in power-law scaling exponents of ligation frequencies versus genomic distance. In the limit of low ligation rates and high digestion efficiencies (for fixed crosslinker concentrations), structural degradation of Hi-C maps ensues from significant displacement of cleaved DNA fragments away from their initial positions in the original chromosome conformations of the native ensemble. A tendency towards the flattening of the power-law scalings of ligation frequencies results from information loss in ligation maps, compared to contact maps of the native ensemble of structures. Increasing crosslinker concentration fosters genome architecture preservation, power-law exponents of the ligation frequencies closely resembling the native ensemble’s contact probability.

The protocol enables the exploration of effects of numerical algorithms for the post-processing of Hi-C data, such as matrix-balancing methods. Although there is no appreciable visual difference between pre and post KR-balanced maps, normalization depletes count-enriched domains with at the expense of enriching other parts of the map. The effect of KR-normalization, as far as SVD of the PCM of the maps is concerned, is not significant or appreciable relative to the raw data, as revealed by the singular value spectra and FPC for raw and KR-balanced data. Insofar as PCA of the maps is concerned, the effect of matrix balancing is minimal and a hypothesis about uniformity on the average column sum of the maps is consistent with raw data. This suggests that, to the extent that PCA is concerned, the FPC is insensitive to KR-normalization and numerical post-processing via matrix balancing is justified, consistent with a uniform average number of counts per locus with small locus-to-locus fluctuations. However, compartment switches can be detected from column sums in raw contact maps, with measurable enrichments and depletions in the final KR-balanced maps compared to the originals.

In-silico Hi-C provides sheds light on the dependence of ligation probability with respect to Euclidean distance, a relevant question that experimental methods have not addressed thus far. Numerical experiments suggest that this dependence is sigmoidal, with a characteristic distance over which proximity ligations typically occur, on the order of some 20[nm]. Hence, in-silico experiments of proximity ligation assays can numerically estimate the extent of locality of ligation events.

Numerical experiments show that crosslinking DNA to an underlying nuclear protein matrix percolating across the nucleus is consistent with results of actual Hi-C experiments. This idea of the matrix, entertained in the literature since the 1970s and about which a measure of controversy and discussion have revolved, was revived upon arrival of recent experimental evidence pointing towards its actual existence.^42–48^ Consistency between in-silico results in this work, assuming the existence of the matrix, and actual Hi-C experiments does not rule out other hypotheses for the way crosslinking occurs in real laboratory settings. However, it suggests that DNA may possibly crosslink via long-range bridges in a protein network that endows the nucleus with a measure of structural rigidity.

PaleoHi-C successfully retrieved chromosome conformations from a 52,000 year-old woolly mammoth’s skin cells. An explanation for the experiment’s success suggests a phase transition of the nuclear material to a glassy state (“*chromoglass*”), given the dryness and low temperatures of the Siberian permafrost, enabling chromosome fossilization.^36^ If protein matrices are actual structural features of nuclei, perhaps an alternative explanation for the exquisite preservation of the woolly-mammoth’s three-dimensional genome architecture could involve the matrix’s potential role in crosslinking DNA oligonucleotides, enabling the survival of chromosome conformations in ancient samples after tens of thousands of years.

## 6. ACKNOWLEDGEMENTS

We thank the members of the Nuclear Physics working group in the Center for Theoretical Biological Physics for valuable discussion. This research was supported by the National Science Foundation through the awards PHY-2412651 and PHY-2019745, and by the National Institute of General Medical Sciences of the NIH under award R35-GM146852. The content is solely the responsibility of the authors and does not necessarily represent the official views of the funding agencies. We are grateful to Patrick Fodor for administrative help.

## 7. AUTHOR CONTRIBUTIONS

In this work, all authors contributed with the writing of the manuscript. M.D.P., B.J.Z., and A.D. conceived the project. B.J.Z., and M.D.P. led the development of the concepts. B.J.Z., A.D. and L.B. performed the simulations and prepared the figures. L.B. provided the statistical analysis about restriction sites for enzymatic digestion on actual genome sequences. A.W. provided the ensemble of structures for the simulations. All authors contributed to the final version of this work.

## 8. COMPETING INTERESTS

The authors declare no competing interests.

## 9. DATA AND CODE AVAILIBILITY

**Ensemble of native structures used in in-silico Hi-C simulations.** An ensemble comprised of 5000 native structures representative of a 1.1Mbp section (genomic region: 95.4 to 96.5 Mbp) of human lymphoblastoid cells at a 200bp resolution was generated from NuChrom, a nucleosome resolution model for Hi-C map inversion trained on experimental ensemble maps.^50^

Code for this model is available at https://github.com/DiPierroLab/NuChroM.

**In-silico simulator of Hi-C experiments.** The code developed in this project for the numerical implementation of the in-silico protocol for proximity ligation assays includes the effective modeling of crosslinking, digestion with restriction enzymes and ligations, using the molecular dynamics Python library HOOMD-blue.^54^ In addition, code that implements the FIRE algorithm at a preparatory/pre-processing stage, as well as post-processing code for aggregation of ligations, construction of ligation and contact maps, and calculations of ligation frequencies versus genomic and Euclidean distances was also developed. All these codes are freely available at https://github.com/DiPierroLab/Bernardo.

**Knight-Ruiz (KR) algorithm for matrix balancing.** A numerical implementation of the Knight-Ruiz algorithm was borrowed for the purposes of balancing the in-silico Hi-C maps from the simulations. In particular, the KR code used in this work is included as a function in the freely available Hi-C Matrix Balancing (HCMB) code.^55^ The original Python code for the KR algorithm function contained in HCMB was developed in the context of gcMapExplorer.^56^ The KR function of the HCMB code can be freely accessed at https://github.com/HUST-DataMan/HCMB.

**Statistical analysis of restriction enzyme digestion sites.** The human genetic sequences analyzed in the context of the study of restriction site statistics are freely available from the Telomere-to-Telomere Consortium. In particular, in this work we used the T2T-CHM13 reference genome.^57^ The code developed for the analysis of restriction site density and the distribution of distances between consecutive restriction sites in the human genome is freely available at https://github.com/lrburack/restriction-site-distribution.

## 10. METHODS

### Details on the molecular dynamics of the in-silico protocol for numerical proximity ligation assay experiments

As was sketched in Section 2 of the main body of this work, the numerical experiments of Hi-C involve the effective computational representation of crosslinking, digestion and ligation reactions. The simulation of these steps by means of molecular dynamics involves the specification of a homopolymer Hamiltonian U_HP_ dependent on the collection of positions 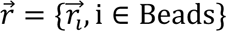 of all the nucleosomes (beads) in the structures. The terms in the Hamiltonian include shifted FENE bonds, truncated Lennard-Jones potentials for hard-core repulsion (volume exclusion) and harmonic angle potentials

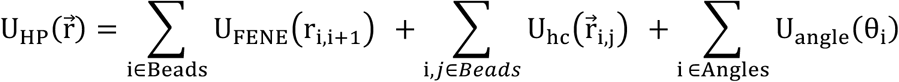

where *Beads* is the set of all beads (nucleosomes), *Angles* is the set of all angles in the polymer, and the aforementioned potentials and bonds are defined as follows:

- Shifted FENE (Finitely Extensible Nonlinear Elastic) bonds *U_FENE_* are defined as:

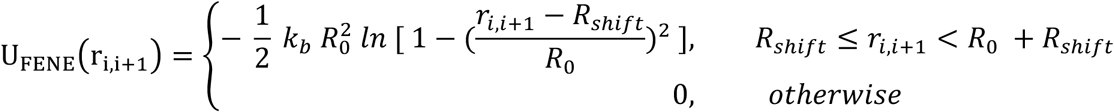

with 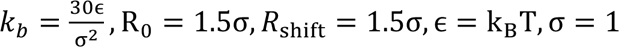 (10 [nm] in real size).
- Hard-core repulsive potentials *U_hc_* between pairs of beads are defined by means of truncated Lennard-Jones interactions:

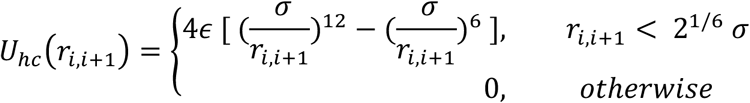
- Harmonic angle potentials *U_angle_* are defined as:

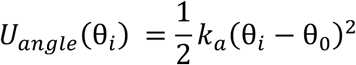

with k_a_ = 4ɛ and 𝜃_0_ = 𝜋.

## Notes

### Competing Interest Statement

The authors have declared no competing interest.

