## Supplementary Information for "Understanding the physical processes behind DNA-DNA proximity ligation assays"

**The digestion phase of Hi-C can be treated as a uniform random process**

The in-silico Hi-C protocol used in this work assumes that the digestion phase of the Hi-C experiment is a uniform random process. That is, all bonds between beads have equal probability of being digested. To assess the validity of this assumption, we searched the T2T-CHM13 reference genome (from the Telomere-to-Telomere Consortium) for restriction sites of three restriction enzymes: MboI (GATC), HindIII (AAGCTT), NcoI (CCATGG).[1] Restriction sites of the analyzed enzymes were found to be uniformly distributed in the human genome, as suggested by Figures S1 and S2.

Figure S1 shows the number of restriction sites found in bins of genomic regions that are 1Mbp long. We observed regions with vastly different restriction site density. These mostly lie either at the centromere region or at the end of the chromosome (telomeres) and were excluded from our analysis. The regions around the centromeres (chr1: 115-150Mbp; chr7: 50-70Mbp) were removed, as well as the first and last 20Mbp of each chromosome were also removed. The restriction site density analysis on the remaining genomic regions shows roughly constant signals for the three enzyme restriction sequences considered, with some measure of fluctuation around the mean. Whilst there are fluctuations that show as a few significant sharp peaks and troughs around the mean (showing isolated regions of sharp enrichment or depletion), the overall pattern suggests a relatively stable mean about which said fluctuations take place. This is roughly consistent with Poisson process statistics, for which the mean number of uncorrelated events in an interval of a given length is fixed. Therefore, it is roughly consistent with an ansatz of uniformly distributed restriction sites along the genome.

Further statistical analysis on restriction site data for the 3 enzymes is presented in Figure S2, which shows the probability distributions for the genomic distances between consecutive restriction sites along the genome. If restriction sites are uniformly distributed across the genome in an uncorrelated way, the consecutive distances between restriction sites should be exponentially distributed. Given a restriction enzyme motif with the same probability $p_m$ of appearing anywhere in the genome, the probability of $x$ consecutive bases containing no restriction sites is $p(x) = (1 - p_m)^x \propto a^{-bx}$.

The distribution of genomic distances between consecutive restriction sites was compared to this theoretical prediction. The prediction was normalized with $-\ln(1 - p_m) * \text{binsize}$, and $p_m$ was determined for each chromosome separately according to $p_m = \frac{\text{Number of Sites}}{\text{Chromosome Length}}$. For each restriction enzyme, this simple model fit the data well. Figure S2 shows evidence consistent with the ansatz of exponentially distributed genomic separations between successive restriction sites, with characteristic lengths of ~400bp for the four-cutter and ~3000bp to ~4000bp for the six-cutters.

The evidence shown in Figures S1 and S2 supports the treatment of the digestion phase of the experiment as a random process, where bonds to be cleaved are picked uniformly at random along the nucleosome resolution native structures.

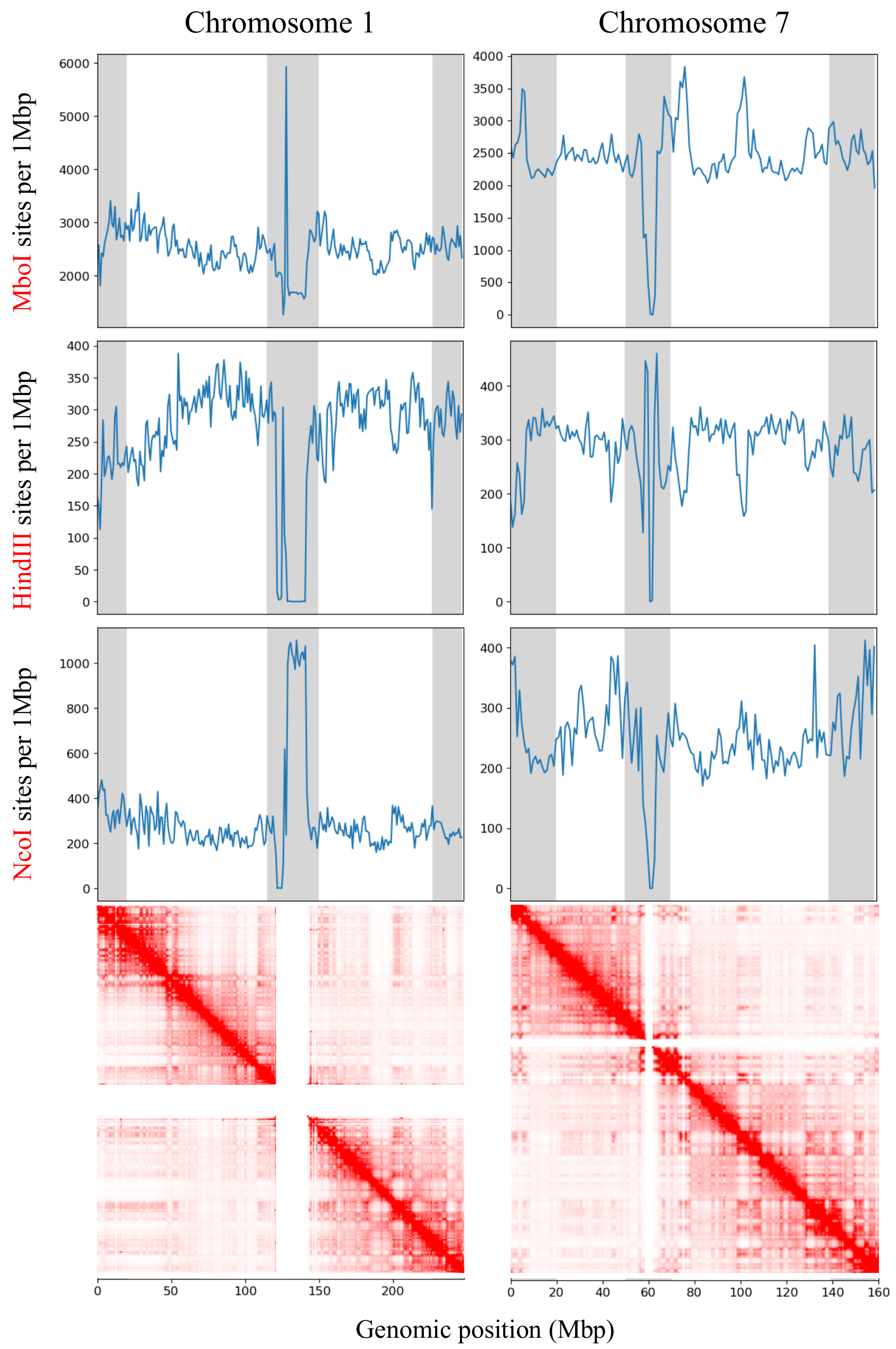


**Figure S1**: **The number of restriction sites in 1Mbp bins for chromosomes 1 and 7.** Regions excluded from further analysis are shown in gray. The corresponding Hi-C maps are shown at the bottom. Regions of anomalous restriction site density were observed near the centromere, where tandem repeat sequences make the region unmappable. The unmappable regions are visible in the corresponding Hi-C maps as thick white stripes, as there is no contact data there.

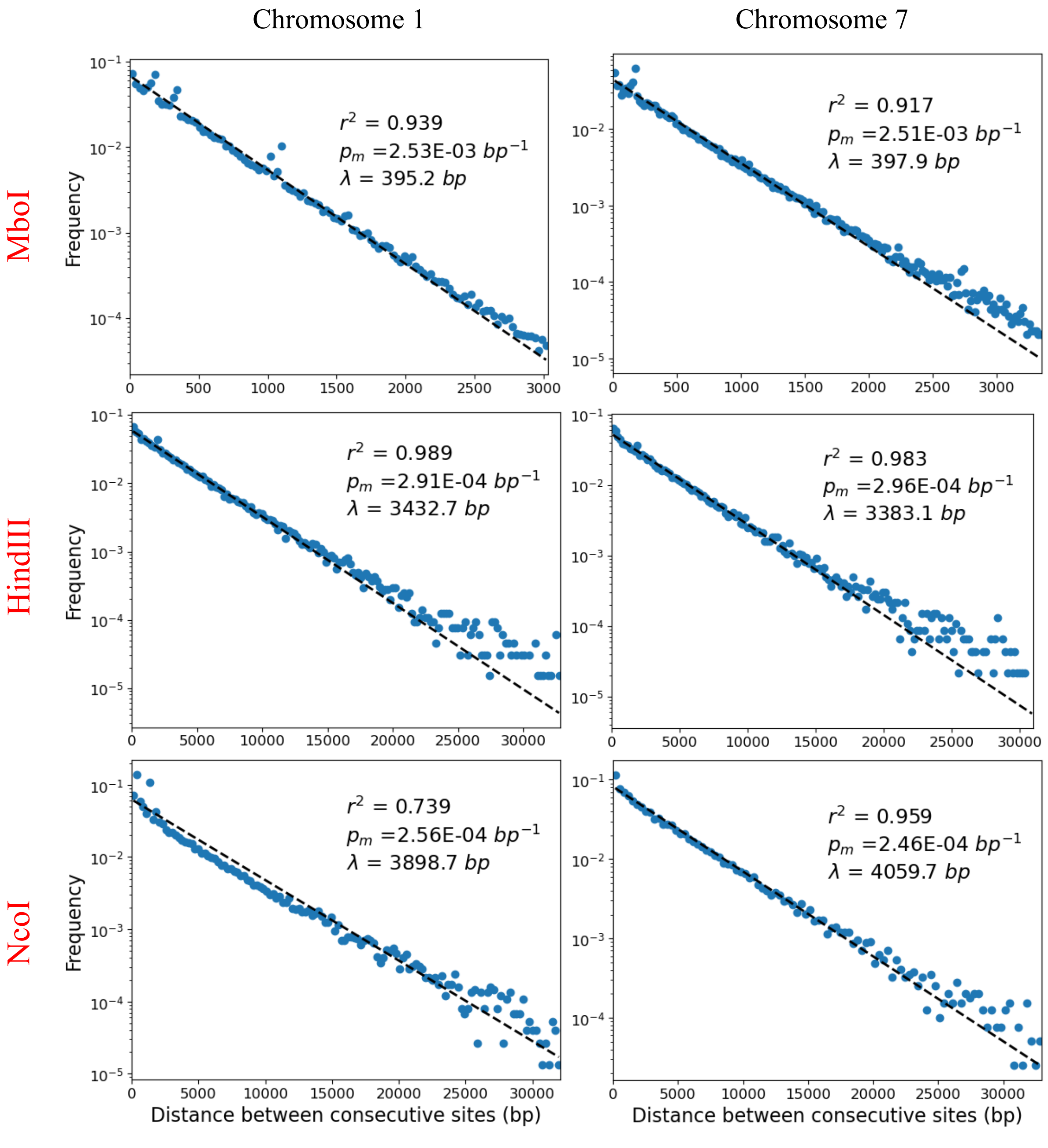


***Figure S2*: The distribution of distances between consecutive restriction sites** (blue scatter), and the model assuming restriction sites are positioned in a uniform random manner (black dashed line). For each chromosome-enzyme pair, the $r^2$ of the model, the enzyme motif probability $p_m$, and the characteristic length $\lambda = \frac{1}{p_m}$ are reported. Some outliers were observed; 99.9% of the distances between consecutive restriction sites are represented above, and the outlying 0.1% were discarded.

**Estimation of timescale for simulations of DNA-DNA proximity ligation experiments.**

To estimate the timescale of the numerical Hi-C experiments, we simulated the motion of a single nucleosome particle under Langevin dynamics. The particle undergoes standard diffusion in the absence of excluded volume effects and bonds. Fitting the mean-squared-displacement (MSD) as a function of the time frame index, one can read the timescale between successive frames with a least-squares fit of the slope of the curve. A nucleosome core particle is modeled as a sphere with a diameter of $d = 11[nm]$ and the nucleus is modeled as an aqueous medium, so that the viscosity is taken to be that of water $\eta \approx 0.89[mPa.s]$ at room temperature $T = 298[K]$. Therefore, as per the Stokes-Einstein equation, the diffusion constant for the nucleosome particle is estimated to be:

$$D = \frac{k_B T}{6\pi\eta r} = \frac{\left(1.38 \times 10^{-23}\left[\frac{J}{K}\right]\right).(298[K])}{6\pi \times (0.89 \times 10^{-3}[Pa.s]).\left(\frac{11}{2} \times 10^{-9}[nm]\right)} = 4.46 \times 10^{7}[nm^2/s]$$

As a consequence of uncorrelated, random fluctuations of Brownian motion, the MSD of the nucleosome scales linearly with time according to $\langle \vec{x}^2 \rangle = 2n_d Dt$, where $\vec{x}$ is the position of the particle, $n_d = 3$ is the number of dimensions of space and $t$ is the physical time elapsed, which is an integral multiple of the timescale $\tau$ between successive frames $t = n\tau$, where $n \in \mathbb{N}$ labels the discrete time-frame index. Fitting the MSD $\langle \vec{x}^2 \rangle$ versus the time-frame index $n$ with a least-squares regression and using the value of the diffusion constant estimated above, we estimate the value of the timescale by measuring the slope of the linear scaling to be $\tau = 2.2678 \pm 0.0008\ [\mu s]$

The result of fitting the MSD versus the time-frame index, averaged over 5500 different simulations of the diffusive process is shown in Figure S3, where the MSD data from the numerical experiments of diffusion is represented with light blue points and the dark blue line represents the least-squares fit to said data.

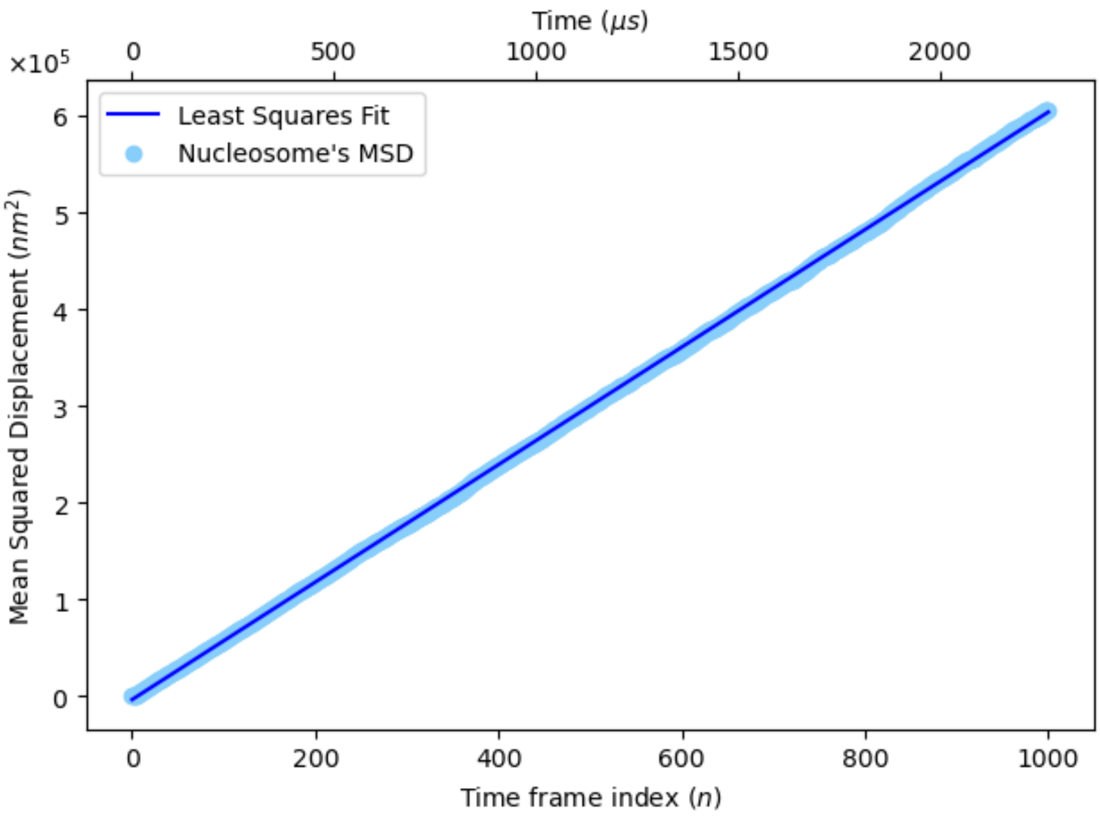


***Figure S3. Mean squared displacement of nucleosome core particle undergoing Brownian motion as a function of the time frame index and physical time.*** The MSD data from the numerical experiments of diffusion, averaged over 5500 independent realizations, is shown with light blue data points as function of the discrete time frame index (lower horizontal axis) and as a function of the estimated physical time scale in microseconds (upper horizontal axis). The dark blue curve is the result of a least-squares regression of the MSD data versus the discrete time frame index to a linear function. The slope the linear fit enables an estimation of the timescale $\tau$ between successive frames by way of the Stokes-Einstein equation.

**Non-equilibrium effects of enzymatic digestion and diffusion on distance maps, contact maps, contact probabilities and their power-law exponents in the absence of crosslinking.**

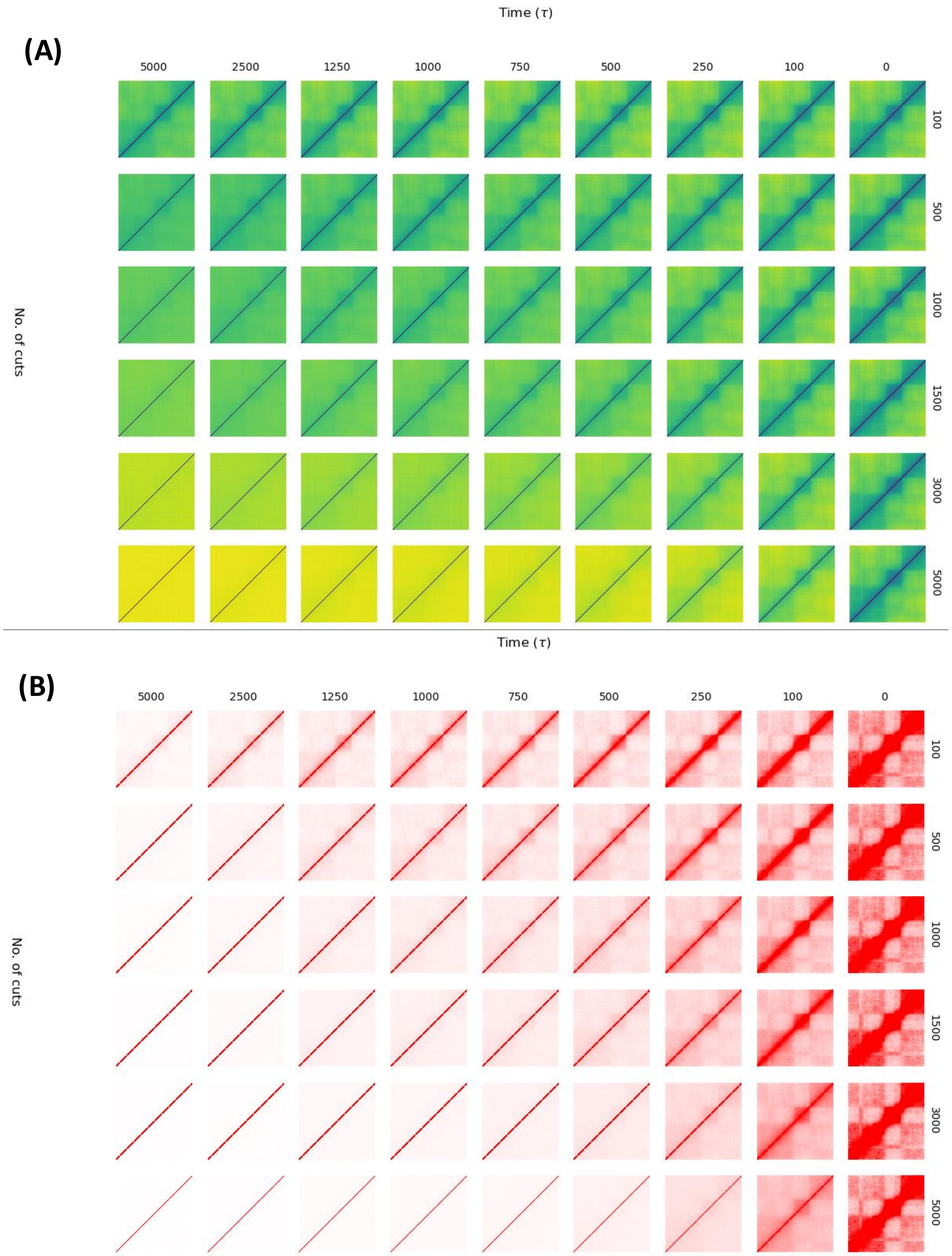


***Figure S4. Non-equilibrium effects of digestion and diffusion on average Euclidean distance maps and average contact maps in the absence of crosslinkers.*** The effects of enzymatic digestion of chromatin into a fragmented structure and the ensuing diffusive motions of these chromatin fragments in the absence of crosslinking agents is assessed by computing the average Euclidean distance maps and the average contact maps over the ensemble of fragmented structures at different stages throughout the time evolution of the out-of-equilibrium expansion. The loss of structure in average distance and average contact maps becomes more pronounced, as expected, for higher number of digested bonds. In the limit when the number of digested fragment ends is such that the chromosome is effectively split into a "gas" of nucleosomes, any discernible structure in the maps is effaced in the long-time limit, as the maps approach constant matrices. ***(A) Average Euclidean distance maps of degraded structures as a function of time and number of digested bonds.*** The grid of average distance maps shows the progressive loss of chromosome conformation structure (as captured by maps) with increasing diffusion and digestion efficiency. The non-equilibrium effect of diffusive motion in time for a fixed number of digested bonds is shown along the rows, since each row corresponds to a different number of initial enzymatic bond cuts. The effect of the number of digested bonds can be assessed along the columns, since each column corresponds to a different point in time in the diffusive process (measured in units of $\tau$). In the limit when all bonds are cut, a nucleosome "gas" obtains, and, in the long-time limit, the loss of discernible structure in the map is so pronounced that it becomes a uniform matrix, all semblance of domains and compartments having been washed away, as evidenced by the flattening effect of the contact probabilities. ***(B) Average contact maps of digested structures as a function of time and number of number of bonds digested.*** Consistent with the picture presented in Fig S4.(A), we show a grid of the average contact maps of the degraded structures, where time (measured in units of $\tau$) changes along the horizontal direction and the initial number of digestion cuts changes along the vertical direction. For a given degraded structure, a contact between two loci exists if said loci find themselves in proximity, separated by a distance of $r = 1.5\sigma = 15\ [nm]$. The loss of structure manifests as a decrease in the number of contacts and the corresponding washing away of any evidence of domains and compartments of the conformations of the native ensembles.

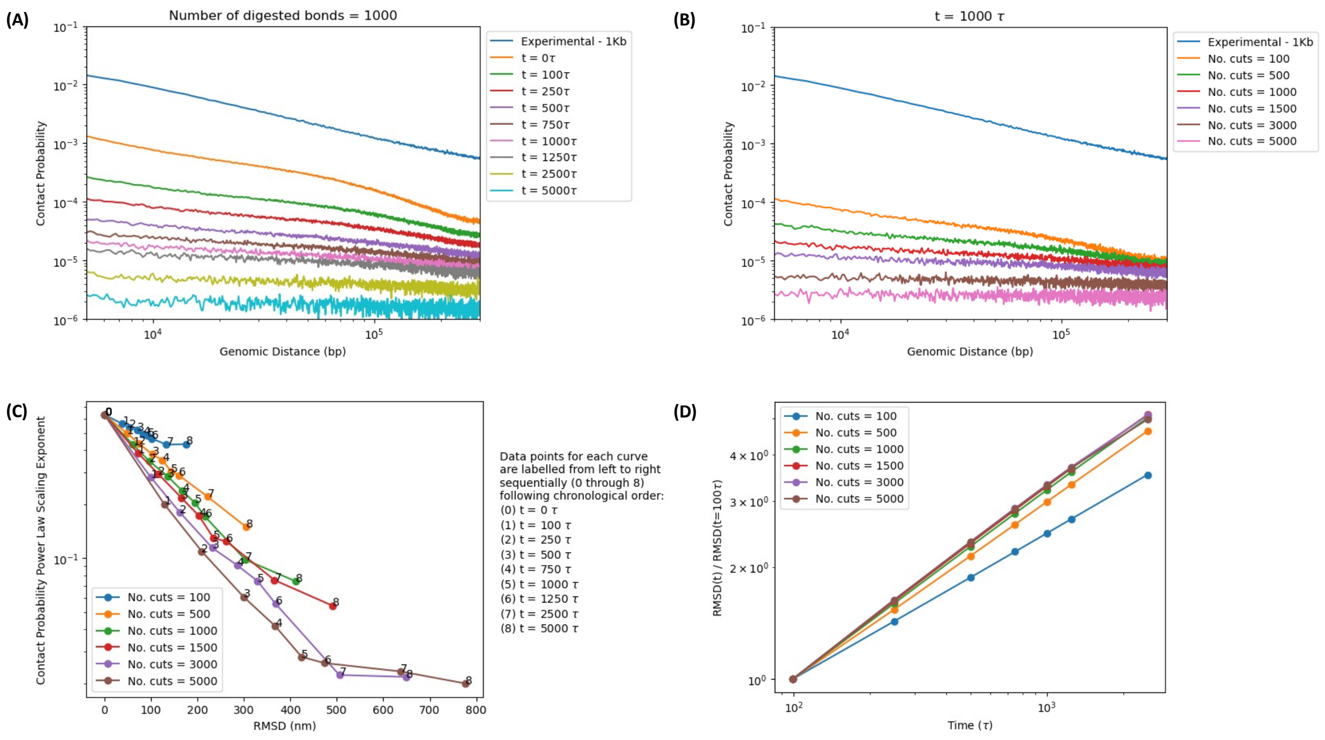


***Figure S5. Non-equilibrium effects of digestion and diffusion on contact probability power law scaling in the absence of crosslinkers.*** The power-law scaling exponent of the contact probability as a function of genomic distance decays with increasing diffusion (and time) for a fixed number of initial digestion cuts. Also, the loss of structure in contact probability becomes more pronounced, as expected, for higher number of digested bonds. In the limit when the number of digested fragment ends is such that the chromosome is split into a "gas" of nucleosomes, the power law exponent tends to zero in the long-time limit and the contact probability flattens, reflecting a complete loss of structure, as was seen in Fig. S4 with progressive effacement of structure tending towards constant distance maps and contact maps. ***(A) Contact probability versus genomic distance at different points throughout time, for a fixed number of enzymatically digested bonds.*** Consistent with Fig. S4, the decay in discernible structure with increasing time (measured in units of $\tau$) for a fixed digestion efficiency is such that the contact probability as a function of genomic distance retains its characteristic power law scaling behavior, but the exponent decreases in time as chromatin fragments diffuse away from their initial positions in the native conformations, resulting in a "flattening" effect on the curves in the long-time limit. ***(B) Contact probability versus genomic distance for a different number of enzymatically digested bonds, at a fixed time.*** Consistent with Fig. S4, the decay in discernible structure with increasing linker DNA digestion efficiency is such that the contact probability as a function of genomic distance also retains its characteristic power law scaling behavior, but the exponent decreases with increasing number of bonds digested, as smaller chromatin fragments (under diffusion) retain less information about the native conformations, resulting in a "flattening" effect on the curves. ***(C) Power law exponents of contact probabilities as a function of average RMSD of nucleosomes for different numbers of***

***initial digestion cuts and at different points in time.*** Power law exponent decays with respect to the average RMSD of nucleosomes, for fixed number of initial digested bonds, with increasing diffusion time. With increasing enzymatic digestion efficiency, the decay of the exponent becomes more pronounced. However, for a fixed value of the exponent, there is a finite range of average RMSD that the nucleosomes could have attained, irrespective of the number of initial cuts, so that the diffusion in space is constrained to remain bounded. ***(D) Average RMSD of nucleosomes as a function of time for different numbers of initial digestion cuts.*** The average RMSD of the nucleosomes in the digested and diffusing fragments (relative to a particular point in time) are shown to increase as time evolves, as expected for this irreversible and out of equilibrium process. It is clear from the plots that, with increasing number of digested bonds, the average length of chromatin fragments decreases, and the dynamics of their motion tend toward standard diffusion. However, in the limit of low enzymatic efficiency, the average lengths of chromatin fragments are larger and the dynamics of their motion show evidence of anomalous diffusion (sub-diffusivity), as can be clearly seen by the different exponents (slopes) of the curves.

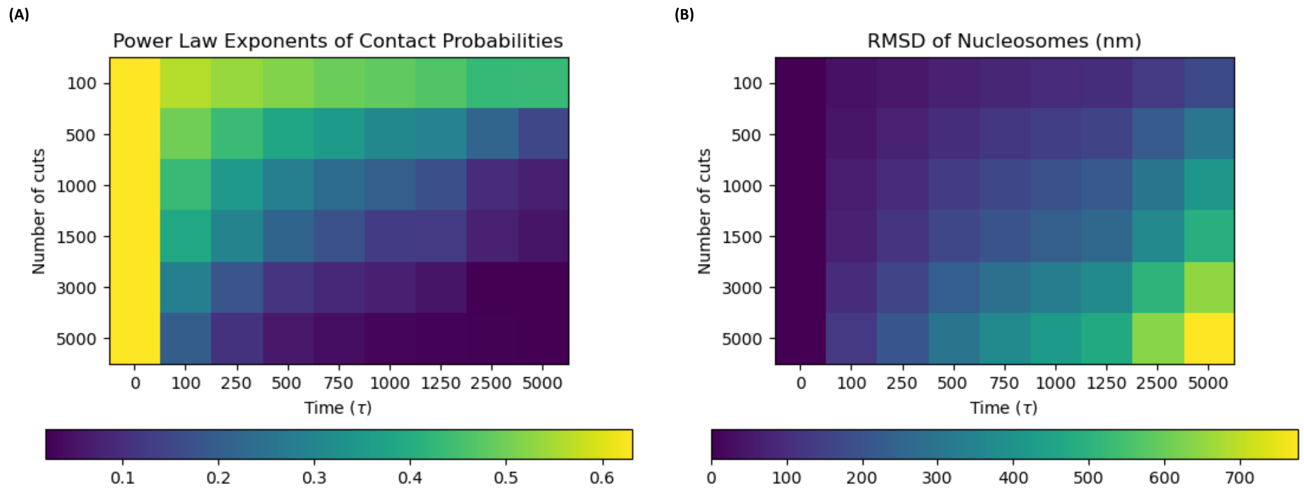


***Figure S6. Non-equilibrium effects of digestion and diffusion on contact probability power law scaling and RMSD of nucleosomes in the absence of crosslinkers.*** Consistent with the results of Figs. S4 and S5, the power-law scaling exponent of the contact probability as a function of genomic distance decays with increasing diffusion and time for a fixed number of initial digestion cuts. Increasing the digestive efficiency of the restriction enzymes or endonucleases favors loss of structure and the flattening of the power law scalings of the contact probabilities. ***(A) Effect of digestive efficiency of enzymes and diffusion time on power law scaling exponents of contact probability.*** The degradation of the power law scaling with increasing diffusion time is relatively well contained for low digestive efficiencies, since chromatin fragments are larger and bulkier and retain more structure of the native conformations, as can be seen for the case with just 100 bond cuts, wheareas, in the nucleosome gas limit of 5000 bond cuts, the flattening decay of the power law in time is very pronounced and any semblance of structure is lost very quickly. ***(B) Effect of digestive efficiency of enzymes and diffusion time on average RMSD of nucleosomes.*** With increasing digestive efficiency, the restriction enzymes or endonucleases are capable of producing smaller and lighter chromatin fragments, whose dynamics will tend to normal diffusion in the limit of the nucleosome gas and higher average RMSDs in the long run, leading to constant contact maps. For smaller number of bonds cut, fragment sizes are larger and heavier and their motions approach anomalous diffusion (subdiffusivity), which is reflected in smaller average RMSDs, consistent with distance maps that still preserve some structure even in the long-time limit.

**Effect of crosslinking efficiencies and fragment diffusion on distance maps, contact maps, contact probabilities and their power-law exponents (with a fixed 1000 digested bonds per structure)**

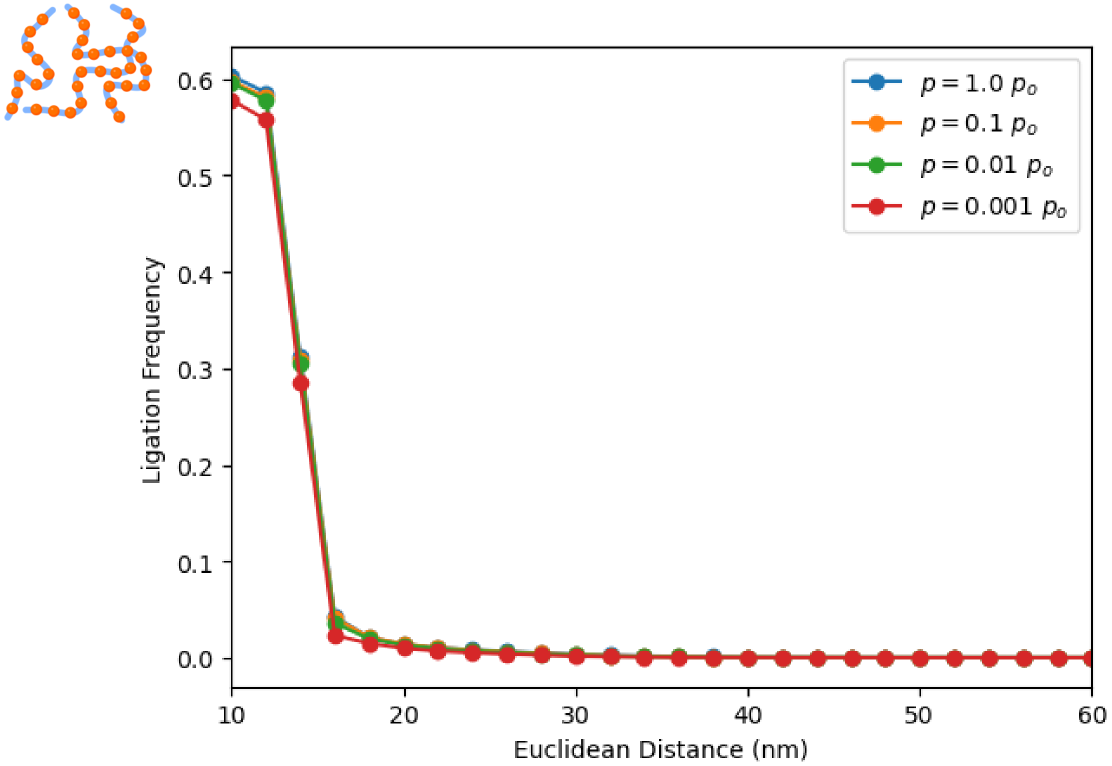


***Figure S7. Ligation frequency as a function of Euclidean distance for high crosslinking efficiency and different ligation rates.*** The ligation frequency as a function of Euclidean distance exhibits the characteristic sigmoidal shape with a very sharp decay that suggests a typical length scale over which ligations take place in the limit of high crosslinking efficiency (5000 nucleosomes crosslinked, i.e., 90.9% of the DNA crosslinked to the protein matrix). This typical length scale is of the order of ~15 [nm], irrespective of the ligation rates. Since the structures are effectively frozen in place and resemble their original native states, the curves collapse on top of one another and non-equilibrium effects due to diffusion are immaterial. Information about the original conformations as captured by the average contact map are very well preserved, since the ligation map very closely resembles the average contact map in this limit. Created in BioRender. Zubillaga, B. (2025) https://BioRender.com/m5gat3n

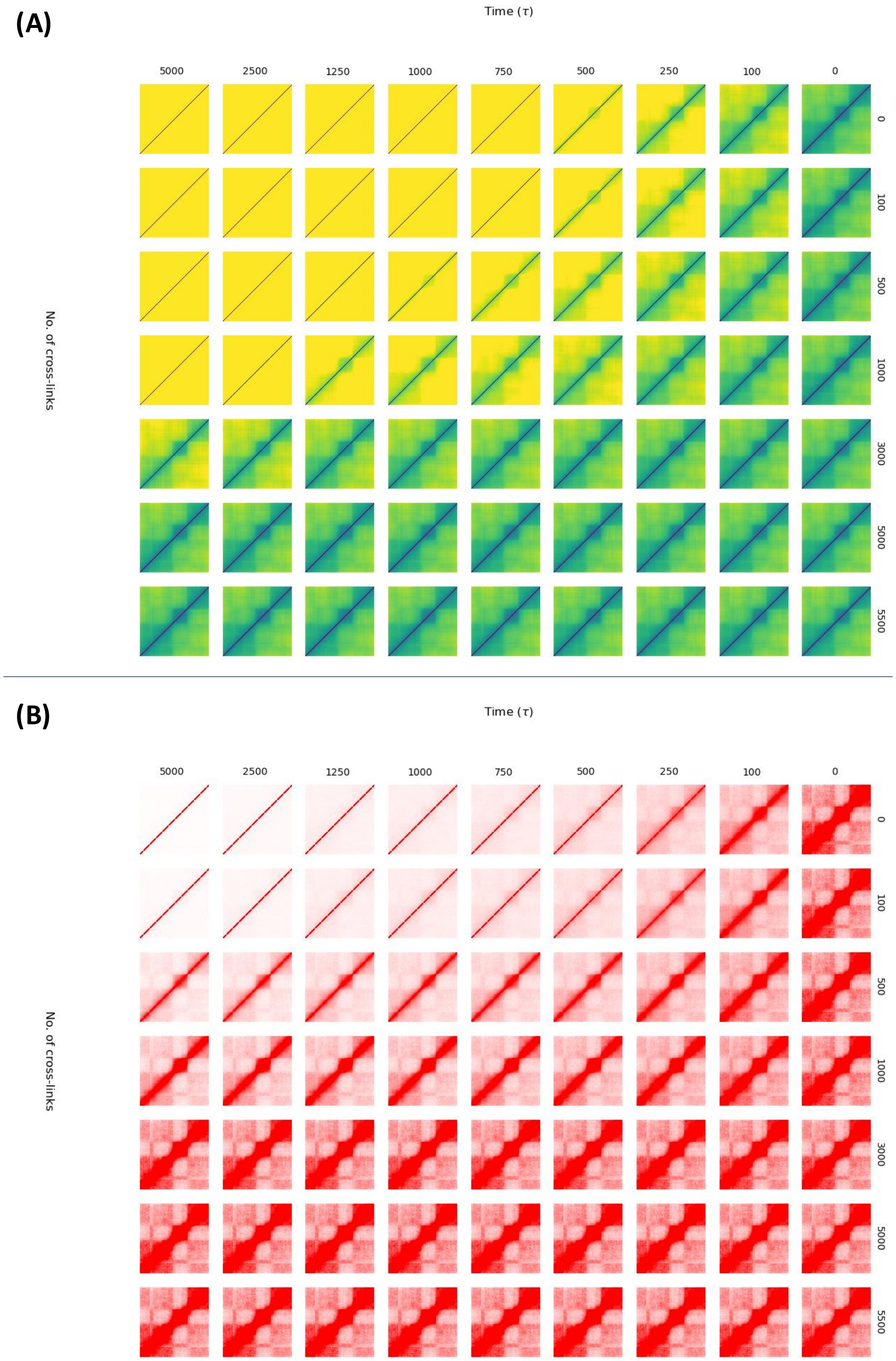


***Figure S8. Effects of crosslinking and diffusion on average Euclidean distance maps and average contact maps.*** The effects of crosslinker concentration, in affixing DNA fragments to the nuclear protein matrix for a fixed enzymatic digestion efficiency is by the average Euclidean distance maps and average contact maps over the ensemble of fragmented structures at different stages throughout the time evolution of the non-equilibrium dynamics. Loss of structure in average distance and average contact maps becomes more pronounced, as expected, for lower numbers of crosslinked nucleosomes. In the limit of no crosslinking, DNA fragments are free to diffuse away from their initial positions in the native conformations and structure is progressively lost in time. In the limit when all nucleosomes are crosslinked to the matrix, the fragments are frozen and information about the conformation is preserved by the map. ***(A) Average Euclidean distance maps of degraded structures as a function of time and number of crosslinks.*** The grid of average distance maps shows the progressive loss of chromosome conformation structure (as captured by maps) with increasing diffusion and decreasing crosslinking efficiency. The non-equilibrium effect of diffusive motion in time for is shown along the rows, each row corresponding to a different number of crosslinks. The effect of the number of crosslinks can be assessed along the columns, each column corresponding to a different point in time in the diffusive process (measured in units of $\tau$). In the long-time limit with no crosslinking, the loss of discernible structure in the map is so pronounced that it becomes a uniform matrix, all semblance of domains and compartments having been washed away. When all nucleosomes become crosslinked, fragments are frozen in place and the map doesn't change in time. ***(B) Average contact maps of digested structures as a function of time and number of crosslinks.*** Consistent with the picture presented in Fig S7.(A), we show a grid of the average contact maps of the degraded structures, where time (measured in units of $\tau$) changes along the horizontal direction and the initial number crosslinks changes along the vertical direction. For a given degraded structure, a contact between two loci exists if they are in proximity, separated by less than $r = 1.5\sigma = 15\ [nm]$. The loss of structure manifests as a decrease in the number of contacts and the corresponding washing away of any evidence of domains and compartments of the conformations of the native ensembles.

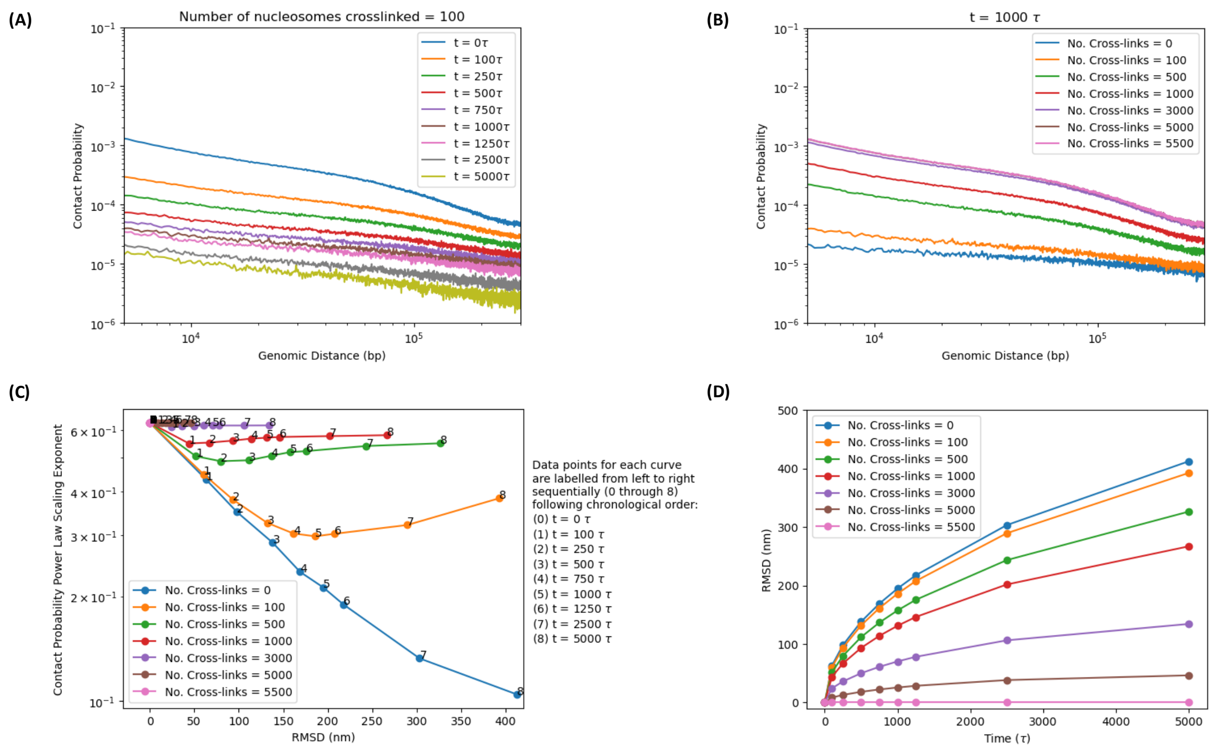


***Figure S9. Effects of crosslinking and diffusion on contact probability power-law scaling.*** The power-law scaling exponent of the contact probability as a function of genomic distance decays with increasing diffusion (and time) for a fixed number of 1000 digested bonds. Also, the loss of structure in contact probability becomes more pronounced, as expected, for lower numbers of crosslinks. In the limit of low crosslinking concentration, DNA fragment motion is unimpeded by the nuclear protein matrix, favoring fragment diffusion and loss of structural information as captured by the contact map. In the limit of high crosslinking concentration, the relative positions between DNA fragments are preserved in time, allowing for the preservation of chromosome conformation as captured by the contact map. This leads to dependences in contact probability scaling exponents on account of diffusion and crosslinking efficiencies, as was suggested in Fig. S7. ***(A) Contact probability versus genomic distance at different points throughout time, for a fixed number of crosslinked nucleosomes.*** Consistent with Fig. S8, the decay in discernible structure with increasing time (measured in units of $\tau$) for a fixed number of crosslinks is such that the contact probability retains its power-law scaling behavior, the exponent decreasing in time as chromatin fragments diffuse away from their initial positions, resulting in a "flattening" effect on the curves in the long-time limit. ***(B) Contact probability versus genomic distance for a different number crosslinked nucleosomes, at a fixed time.*** Consistent with Fig. S4, the decay in discernible structure with decreasing crosslinker efficiency is such that the contact probability as also retains power-law scaling behavior, the exponent decreases with number of crosslinks, as free fragments can diffuse away (subject to excluded volume), thus leading to loss of chromosome conformation information. ***(C) Power-law exponents of contact probabilities as a function of average RMSD of nucleosomes for different numbers crosslinks and at different points in time.***

Power-law exponent decays with respect to the average RMSD of nucleosomes, for fixed number of crosslinks, with increasing diffusion time. With decreasing crosslinking efficiency, the decay of the exponent becomes more pronounced, as diffusion effaces structure in the map. In the limit of high crosslinking efficiency, the native conformations become frozen and the map preserves information. ***(D) Average RMSD of nucleosomes as a function of time for different numbers of crosslinks.*** The average RMSD of the nucleosomes in the digested and diffusing fragments are shown to increase as time evolves, as expected for this irreversible, non-equilibrium process. With decreasing crosslinking efficiency, the dynamics of fragment motion tend toward diffusion, as fragment motion is unimpeded by the nuclear matrix. For high crosslinking concentration, the protein matrix freezes the fragments in place and structure is preserved.

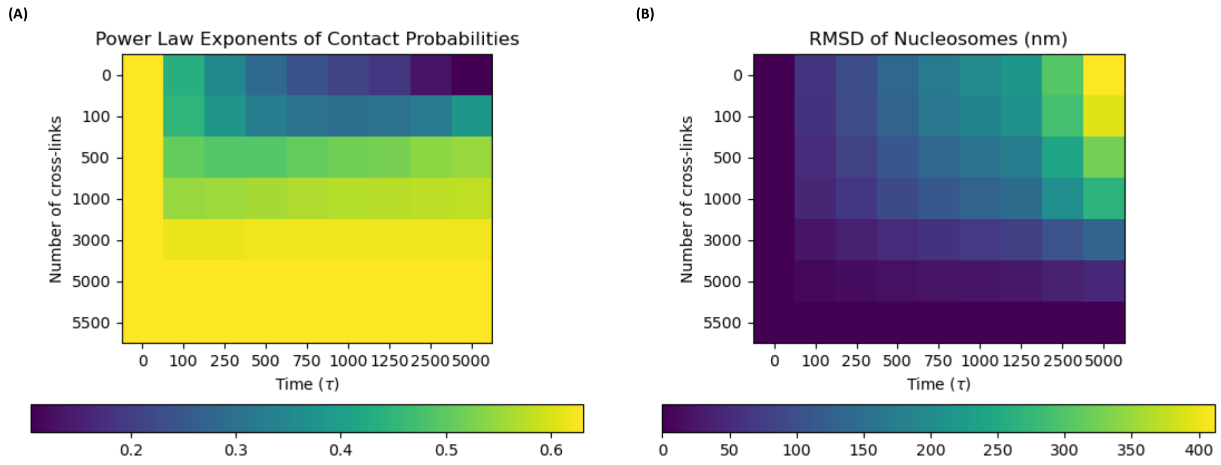


***Figure S10. Effect of crosslinking efficiency and diffusion on contact probability power law scaling and RMSD of nucleosomes.*** Consistent with the results of Figs. S8 and S9, the power-law scaling exponent of the contact probability as a function of genomic distance decays with increasing diffusion and time for a fixed number of crosslinked nucleosomes. Decreasing the number of crosslinks favors loss of structure and the flattening of the power law scalings of the contact probabilities. ***(A) Effect of crosslinking efficiency and diffusion time on power law scaling exponents of contact probability.*** The degradation of the power-law scaling with increasing diffusion time is relatively well contained for high crosslinking concentrations, since chromatin fragments are pinned down to the nuclear protein matrix and thus frozen in place, whereas, in the absence of crosslinking agents, the flattening decay of the power law in time is very pronounced and any semblance of structure is lost very quickly as DNA fragments diffuse away unimpeded. ***(B) Effect of crosslinking efficiency and diffusion time on average RMSD of nucleosomes.*** With decreasing crosslinker concentration, DNA fragments motion is less hampered by the nuclear matrix and their dynamics will tend to normal diffusion, with higher average RMSDs in the long run, leading to constant contact maps. For high crosslinking concentrations, fragments are frozen in place on account of the nuclear matrix, yielding smaller average RMSDs, consistent with distance maps that still preserve structure in the long-time limit.

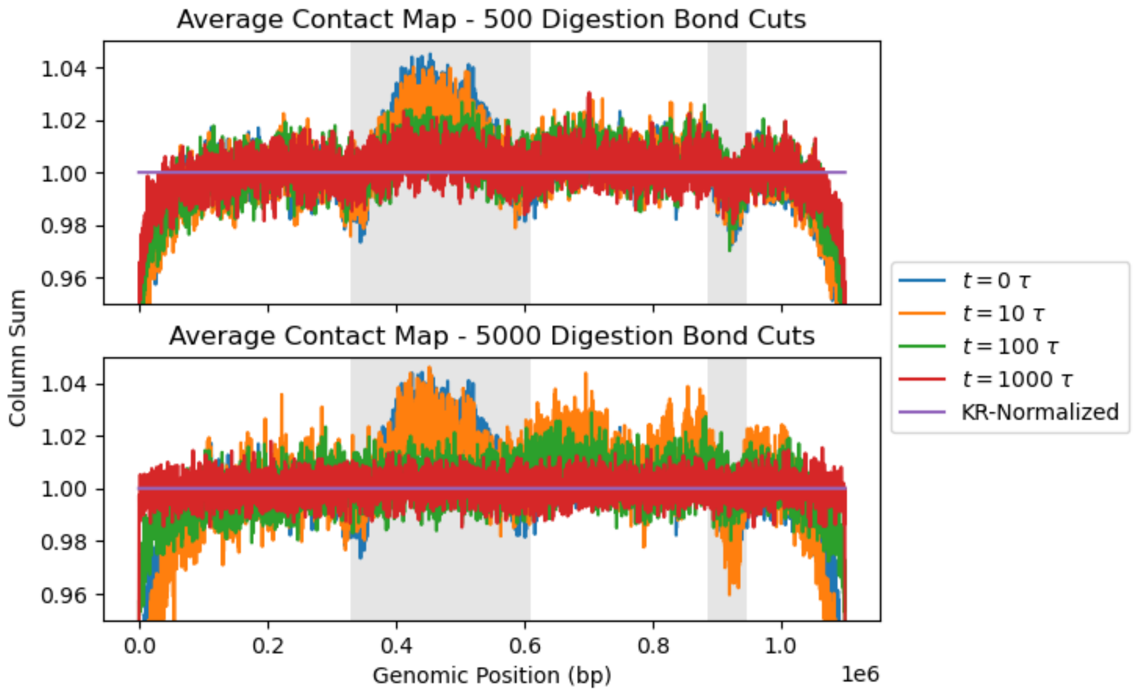


***Figure S10. Non-equilibrium effects of subdiffusive motion of DNA fragments on column sums of the average contact probability as a function of time, for numerical experiments with different digestion efficiencies.*** The upper and lower plots show the column sums of the average contact maps on the ensemble of structures along genomic position for numerical experiments with 500 and 5000 enzymatic bond cuts, resp., at different times throughout the diffusion process. The times at which the snapshots of the ensemble of trajectories of the crosslinked and digested structures were interrogated through contact maps follow in geometric progression, starting at the beginning of the experiment ($t = 0$), when the chromosome conformations are still intact. The effect of diffusion on the degradation of the information of the native conformations captured by the maps is evident from the curves. The central TAD (between genomic positions ~0.3Mbp and ~0.6Mbp), which shows a visible enrichment of contacts (the central "bump" on the gray background) at the beginning of the numerical experiment, is progressively lost as time evolves and DNA fragments are allowed to diffuse away. Likewise, the small depletion on the original ensemble of chromosome conformations around genomic position ~0.9Mbp (shown to the right on gray background) is also progressively lost on account of the non-equilibrium motion of DNA fragments. Comparing the upper and lower plots, one can see that, in the long-time limit ($\mathrm{t} = 1000\tau$), the column sum for the enriched region corresponding to the central TAD is completely effaced in the case of high digestion efficiency (lower plot), but there is still some semblance of enrichment left in the case low digestion efficiency. Likewise, in the long-time limit ($\mathrm{t} = 1000\tau$), the column sum for the depleted region around ~0.9Mbp is completely effaced in the case of high digestion efficiency (lower plot), but there is still some semblance of depletion left in the case low digestion efficiency. This can be understood in light of the fact that, for low digestive efficiency, 500 digested bonds lead to DNA fragments of some ~11 nucleosomes in length (~2200bp) on average, whilst, for high digestion efficiency, 5000 digested bonds lead to DNA fragments of ~1 nucleosome in length (~220bp) on average, i.e., the nucleosome "gas" limit. Also shown, for reference, is the KR-normalized column sum along genomic position, showing the effect of matrix-balancing on the elimination of both fluctuations and enrichments and depletions due to compartment switching.
